# Bacteroidales T6SS minor Hcp subunits form heteromers recognising effectors

**DOI:** 10.1101/2025.01.09.632221

**Authors:** Sergio G. San-Miguel, Manal Kamal Saleh Al-Ammari, Emma Johansson, Anna Kowalska, Uwe Sauer, Paulo Ricardo Batista, Linda Sandblad, Bernt Eric Uhlin, David A. Cisneros

**Affiliations:** Department of Molecular Biology, Umeå University, Sweden; Department of Chemistry, Umeå University, Sweden; Umeå Centre for Microbial Research (UCMR), Umeå University, Sweden; Programa de Computação Científica, Vice-Presidência de Educação Informação e Comunicação, Fundação Oswaldo Cruz, 21040-900 Rio de Janeiro, Brasil; The Laboratory for Molecular Infection Medicine Sweden (MIMS), Umeå University, Sweden; Wellcome-Wolfson Institute for Experimental Medicine, School of Medicine, Dentistry and Biomedical Sciences, Queen’s University Belfast, United Kingdom

## Abstract

The type VI secretion system (T6SS) is a macromolecular protein complex found in Gram-negative bacteria that mediates intercellular antagonism. T6SSs are gut colonization factors that influence gut biodiversity. Hemolysin-coregulated proteins (Hcp) are major structural proteins in these systems. Hcp forms hexameric rings that stack to create an inner tube structure essential for translocating effector proteins into target cells. In Bacteroidales, T6SS loci encode multiple Hcp proteins with unknown function. The gut commensal *Bacteroides fragilis* encodes five Hcp subunits (sHcp and Hcp1-4) that have low sequence similarity. In this study, we investigated the roles of these proteins. Interaction studies showed that sHcp forms homohexamers, which is consistent with a major role of forming the bulk of the inner tube. In contrast, the less abundant minor Hcp1-4 were shown to form an interaction network involving heteromeric complexes. Biochemical analyses demonstrated that Hcp1 and Hcp2 assemble into heterohexamers and that this complex recognizes the secreted effector Bte1. Finally, we showed that Hcp modules, which are encoded in highly syntenic regions in T6SS loci of Bacteroidales, cluster with effectors. These results imply that the minor Hcps genetically cosegregate with cognate effectors, contributing to effector cassette variability. Thus, minor Hcp subunits function as recognition particles for effectors to mediate secretion, which appears to be a conserved trait in Bacteroidales T6SSs. Exploiting these features could facilitate the characterization of unknown effectors by copurifying them with their cognate Hcps. This approach may reveal new insights into bacterial interactions and the mechanisms that establish gut biodiversity.

## INTRODUCTION

The human gut microbiome comprises thousands of species that compete for resources and space. Although the metabolic capability of individual species is a major determinant of stable niche colonization in the gut ^1^, genes that contribute to bacterial antagonism are also well represented in the microbiota (reviewed in ^2^). However, little is known about the molecular mechanisms underlying bacterial antagonism in the gut. Elucidating such mechanisms may contribute to understanding how a biodiverse microbiota is established and how it contributes to intestinal homeostasis.

The type VI secretion system (T6SS) is a macromolecular protein complex encompassing a contractile injection system related to bacteriophages ^3,4^ and forming structures that can span the width of gram-negative bacteria ^3,5^. Soon after its discovery in *Vibrio cholerae* and *Pseudomonas aeruginosa* ^6,7^, it was established that this system participates in bacterial competition by injecting lethal protein effectors. Encoded in T6SS loci are antitoxins, known as immunity proteins, which mediate protection against bacterial kin attacks by neutralizing the activities of effectors ^8^. For bacterial pathogens, T6SSs contribute to disease by injecting antieukaryotic effectors subverting host defences (reviewed in ^9^).

The architecture of the T6SS has mostly been studied in Pseudomonadota and is formed by three main complexes (reviewed recently in ^10–12^). A membrane complex, formed by TssJ, TssL and TssM, anchors the system to the cell envelope. The base plate formed by TssE, TssF, TssG and TssK is attached to the membrane complex. A tubular structure homologous to phage tails, constitutes the contractile complex and is formed by an inner tube complex formed by ring-shaped, hexameric Hcp (TssD) subunits stacked on top of each other, which is enveloped by the contractile tubular sheath formed by TssB and TssC. Sitting atop the inner tube, the VgrG-PAAR tip complex forms a puncturing device, which is housed inside the membrane complex. Upon sheath contraction, the inner tube and tip are injected into target cells. Injected effectors can travel both as attached to the tip complex or as cargo inside Hcp rings ^13,14^. However, the structural and functional features of Hcp-effector complexes are poorly studied.

Gut Bacteroidales include prominent intestinal microbiota species that contribute to polysaccharide degradation in the gut, but individual species such as *Bacteroides fragilis* also participate in gut homeostasis ^15^. T6SS-encoding loci are conserved in gut Bacteroidales and are organized into three highly syntenic genetic architectures (GA): GA1, GA2 and GA3 ^16^. Compared with the better characterised Pseudomonadota T6SSs, Bacteroidales T6SSs diverge in sequence and exhibit important differences ^16,17^. For example, the TssJLM complex is not present and is substituted by another megadalton structure constituted by novel proteins named TssN, TssO, TssP, TssQ, and TssR ^16,18^. Another difference in Bacteroidales, is that effectors are encoded in variable regions within T6SS loci, which differ at the strain level ^16,19^. Consequently, gut Bacteroidales are capable of engaging in interspecific and interstrain competition ^19–21^. Therefore, the T6SS is considered a *bona fide* colonization factor for *Bacteroides* spp. and has been shown to contribute to shaping biodiversity in the gut, especially in infants ^22^.

A conserved feature among Bacteroidales T6SSs, important to this study, is the presence of a multiplicity of Hcp subunits with low sequence similarity in GA2 and GA3 ^16,23^. Although extra *hcp* genes have been observed in other organisms, they are frequently found outside T6SS loci in auxiliary modules ^24–26^. In all Bacteroidales GA2 or GA3 T6SS five Hcp subunits are consistently present, which suggests specific and important roles for these proteins. Interestingly, genes encoding variable effector cassettes are adjacent to these *hcp* genes and bioinformatic analyses suggest that such variability may be specifically regulated ^23^. However, we do not know how multiple Hcp subunits interact or whether they participate in assembly or secretion reactions.

In the present study, we probed the existence of protein‒protein interactions among Bacteroidales Hcp proteins using the T6SS of *Bacteroides fragilis* as a model. Using biochemical approaches, we found that a subset of Hcp subunits, which we define here as minor Hcp subunits on the basis of their lower abundance, establish heteromeric interactions, forming chimeric Hcp particles. These particles were shown to interact with cognate partners, recognising effectors for secretion. Our results demonstrate that the characterisation of numerous Bacteroides T6SS effectors bound to cognate Hcp heteromeres can unlock the study of the molecular mechanisms mediating bacterial competition in the gut microbiota.

## RESULTS

### *B. fragilis* Hcp subunits establish heteromeric interactions

The T6SS locus of *B. fragilis* encodes five Hcp genes: *sHcp* (gene BF9343_1943), *hcp1* (BF9343_1939), *hcp2* (BF9343_1938), *hcp3* (gBF9343_1935), and *hcp4* (BF9343_1934) (Figure 1A). We hypothesised that, similar to other Hcp proteins (reviewed in ^2^), *B. fragilis* Hcps would form ring-shaped hexamers. Therefore, we set out to test their organisation and, specifically, to determine whether the Hcp rings would assemble or stack in a specific order. We performed bacterial two-hybrid assays ^27^ for the five *B. fragilis hcp* genes with N- and C-terminal fusions with the *Bordetella pertussis* T18 and T25 *cyaA* fragments. For each *hcp* gene, four fusion genes were generated, resulting in four possible combinations to test all pairwise interactions (Figure 1B). We cotransformed *Escherichia coli* BTH101 with two plasmids carrying the fusion genes and analysed the appearance of transformants on plates supplemented with X-gal. To define the best parameters at which these interactions occur, we used various IPTG concentrations and growth on plates at 30 °C or 37 °C. To our surprise, only fusions with sHcp showed positive interactions with itself in all four two-hybrid combinations and conditions, suggesting that not all combinations lead to the formation of multimers (Figures 1C and S1). The finding that only this subunit can form homomers, together with the fact that it is the most upstream of the T6SS locus, suggests that, as previously suspected, this gene encodes a structural Hcp protein ^23^. Hcp1-Hcp2 also appeared consistently in all four tested combinations. An interaction between Hcp3 and Hcp4 was detected in most combinations (Figure 1C). Other heterointeractions, such as sHcp-Hcp3, Hcp1-Hcp4 and Hcp2-Hcp3, appeared only in some tested combinations (Figure 1C, magenta, cyan and blue dashed squares, respectively). The Hcp2-Hcp2 interaction occurred in only one combination (Figure 1C, C-term *vs*. N-term). To quantify an average level of interaction between Hcp proteins in all four combinations, we performed densitometric analysis to quantify the appearance of blue hydrolysed X-gal normalized to the positive control (Figures S1 and S2). Averaging across all the tested IPTG concentrations and growth conditions and all four fusion combinations, we obtained a map of interactions, which confirmed our visual observations (Figure 1D). Moreover, the frequency map revealed a hierarchical interaction cascade in which sHcp interacted with Hcp1, which interacted with Hcp2, and so on. Varying the IPTG concentration had little effect on the level of interaction. However, we inexplicably observed that some interactions were seemingly lacking at 30 °C or 37 °C or inhibited bacterial growth (Figures S1, S3D and S3E). Of note, a few interactions were detectable only after prolonged (36 h) incubation (Figures S1 and S2). Taken together, these results suggest that only sHcp has a strong tendency to form homo-oligomeres but that the other Hcps hierarchically form hetero-oligomers.

**Figure 1.**
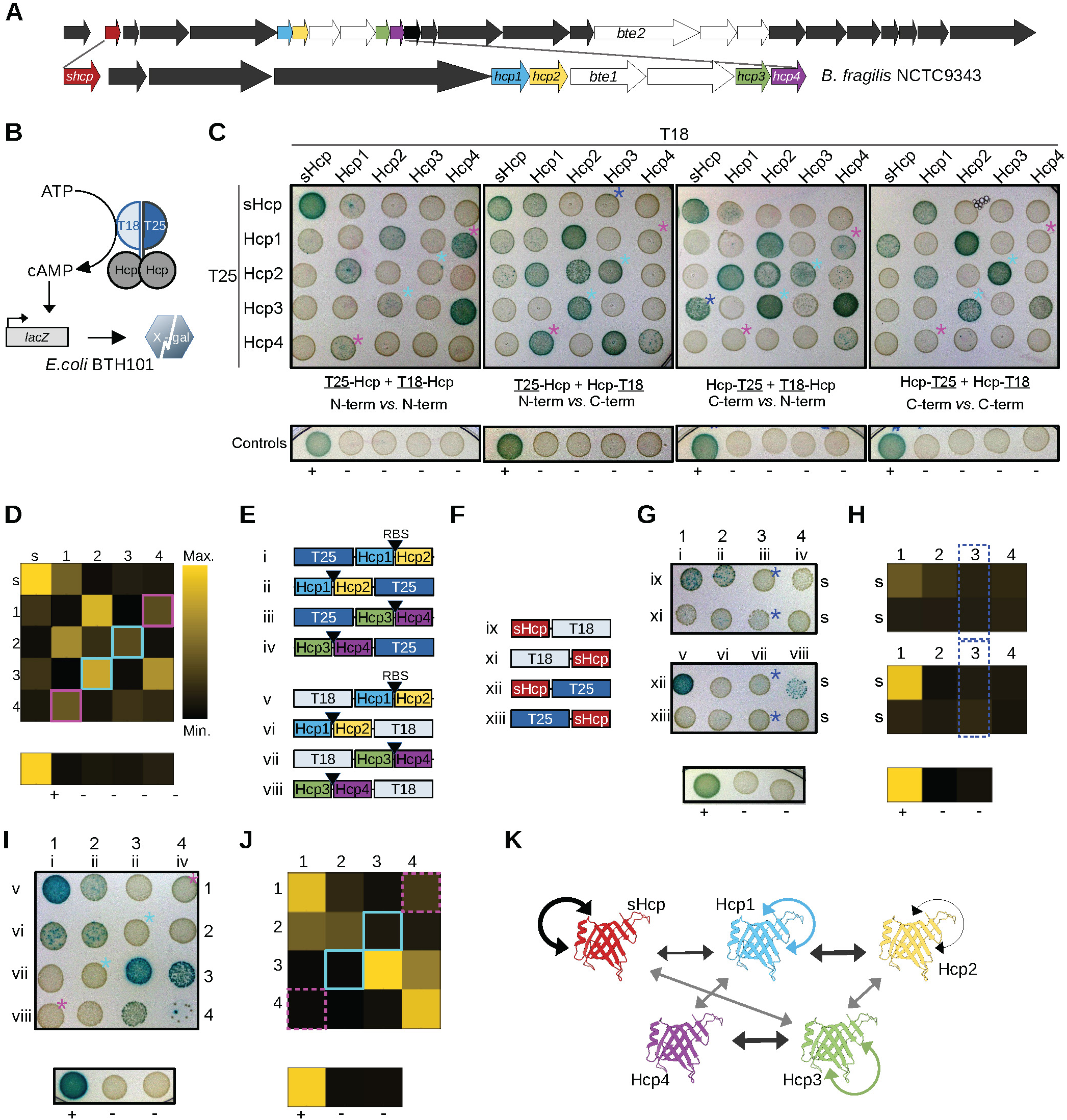
Two-hybrid assays of the major and minor Hcp proteins from *B. fragilis*’ T6SS. **A)** Scheme of *B. fragilis* NCTC9343 T6SS locus. **B)** Scheme of Hcp proteins fused to the *Bordetella pertussis* T18 and T25 two-hybrid fragments in the *cyaA-E. coli* strain BTH101. Interaction of fused proteins leads to the expression of β-galactosidase and hydrolysis of X-gal. **C)** Photographic images of two-hybrid assays with plasmids carrying genes encoding Hcp proteins fused to either a C-terminal T18 and T25 fragments(left panel) or a N-terminal T25 and a C-terminal T18 fragment (centre left panel) or a C-terminal T25 and a N-terminal T18 fragment (centre-right panel) or a N-terminal T18 and T25 fragments (right panel). Asterisks denote interactions inhibited in panels G-J (squares). Hcp proteins are labelled above. The positive control (+) is the dimerization of the yeast GCN4 leucine zipper and the negative controls (-) are the empty vectors in four different combinations. Pictures were taken at 24h from plates containing 250 mM IPTG at 30 °C. **D)** Quantification of two-hybrid assays. The average hydrolized X-gal of all four experiments shown in B, plotted as a color scale in arbitrary units normalized to both the GCN4 leucine zipper (Zip) positive and negative controls. The color scale between black and yellow shows the minimum and maximum, respectively in that plot. The data represents the average of data obtained with varying concentrations of IPTG, at 24h and 36h and at 30 °C and 37 °C. **E)** Scheme of genes coding simultaneous expression of two Hcp proteins fused to T18 and/or T25 fragments used for two-hybrid assays shown in G-J. An artificial *E. coli* ribosome binding site (RBS) was placed upstream of genes encoding a second Hcp subunit. **F)** Scheme of genes coding for N- and C-terminal sHcp proteins fused to the T18 and T25 fragments used for two-hybrid assays shown in G-H. **G)** Photographic images of competitive two-hybrid assay with plasmids encoding sHcp (s) fused to N- or C-terminal T18 or T25 fragments tested for pairwise interactions against T18 or T25 fusions of Hcp1 (1), Hcp2 (2), Hcp3 (3) and Hcp4 (4) in the presence of Hcp2, Hcp1, Hcp4 and Hcp3, respectively. Pictures were taken at 36h. **H)** Quantification of G. **I)** Photographic images of competitive and cooperative two-hybrid assay for pairwise interactions among T18 or T25 translational fusions of Hcp1 (1), Hcp2 (2), Hcp3 (3) and Hcp4 (4) in the presence of Hcp2, Hcp1, Hcp4 and Hcp3, respectively. Pictures were taken at 36h. **J)** Quantification of I **K)** Scheme of interactions among Hcp subunits derived for two-hybrid assays. The wider arrows represent the most frequently observed interactions. Grey arrows represent interactions that are inhibited by other pairwise interactions. Coloured arrows represent cooperative interactions that depend on another interaction.

### Hcp subunits interact through cooperative binding to cognate pairs

Two of the most frequently observed interactions occurred between the products of adjacent genes, namely, Hcp1-Hcp2 and Hcp3-Hcp4. Moreover, there is a highly syntenic arrangement in these two pairs of *hcp* genes across Bacteroides T6SSs with GA2 or GA3, suggesting that the formation of these heterocomplexes has a functional link. Therefore, we aimed to test whether new interactions could be observed when these two pairs of genes with one gene fused to T18 or T25 gene fragments are coexpressed and tested in two hybrid assays against other tagged Hcps. We cloned two contiguous *hcp* genes and inserted an *E. coli* Shine–Dalgarno sequence between them to ensure translation. In this arrangement, only one *hcp* gene is fused to the T18 or T25 fragment (Figure 1E). First, we tested the interactions between sHcp (Figure 1F) and the other Hcps in the presence of their adjacent genes via two-hybrid assays (Figure 1G). We did not find any new interactions for sHcp in this two-hybrid combination. Averaged across the four experimental combinations, only sHcp and Hcp1 were able to interact in the presence of Hcp2 (Figure 1H), as previously observed in the pairwise setting (Figure 1C). However, sHcp-Hcp3, which showed a level of interaction in the pairwise setting, did not interact on average (see blue dashed squares in Figs. 1C and 1H).

We also wanted to determine whether new interactions would occur when adjacent pairs of Hcp proteins were tested via two-hybrid assays. We observed an Hcp1-Hcp1 interaction when T18 and T25 Hcp1 fusions were expressed in the presence of Hcp2 (Figure 1I and 1J), which was not observed in Hcp2-Hcp2 pairwise two-hybrid assays. Similarly, we observed an Hcp3-Hcp3 interaction when the T18 and T25 Hcp3 fusions were expressed in the presence of Hcp4 (Figure 1I and 1J). These results suggest that Hcp1 and Hcp3 can partake in homointeractions but that they exclusively and cooperatively occur in the presence of another Hcp. Our results also showed that some observed heterointeractions are outcompeted by the expression of a second Hcp: specifically, Hcp1-Hcp4 disappears when Hcp3 and Hcp4 are present and Hcp2-Hcp3 disappears when Hcp1 and Hcp2 are produced (Figure 1C, 1I and 1J, cyan and magenta squares). These results suggest that Bacteroidales Hcp proteins form a complex network of interactions (schematic in Figure 1K) but that neighbouring *hcp* genes encode cognate pairs that preferentially interact with each other, so that Hcp1 preferentially binds to Hcp2 and Hcp3 to Hcp4.

### Hcp heterocomplexes form Hcp rings

To obtain structural insight into the *B. fragilis* Hcp proteins, we purified two of the complexes described above. As a control for hexamer formation, we purified a His-tagged version of sHcp (Figure 2A, sHcp-His^6^) which, as observed by negative staining electron microscopy, appeared as rings (Figure 2B). We obtained a crystal structure of sHcp-His^6^ (Figures 2C, S4A and Table S1), which was identical to the VgrG-free apo-structure of He *et al.* ^28^. We subsequently coexpressed Hcp1 and Hcp2-His^6^ for affinity purification, which revealed that Hcp1 readily copurifies with Hcp2 (Figure 2A). Observation of the Hcp1-Hcp2-His^6^ complex via negative staining electron microscopy also revealed the formation of rings that appeared similar to those formed by sHcp (Figure 2B). Cryo-EM observations provided a 2D structure and revealed that the fold of Hcp1-Hcp2-His^6^ was that of canonical Hcps, as shown by the presence of an elongated structure corresponding to β-strands and a circular structure corresponding to the α-helix (Figures 2C, 2D, 2E asterisks and S4B). AlphaFold models of monomeric Hcp1 and Hcp2 (Figures 2F, 2G and S4C) showed a classic Hcp fold. However, a few structural differences were notable. In the case of Hcp1, an insertion between residues G11 and S25 forms a short alpha helix that protrudes from the external beta sheet, and in the case of Hcp2, an N-terminal insertion between residues M1 and L13 is unstructured. In both structures, compared with the sHcp sequence, insertions presented lower PLDDT scores. Purification of the tagged versions of Hcp1 or Hcp2 alone did not yield soluble proteins, which suggests that the interactions between these proteins are necessary for their folding and assembly into rings. These results show that Hcp1 and Hcp2, are binding partners that cooperatively assemble into hexameric rings.

**Figure 2.**
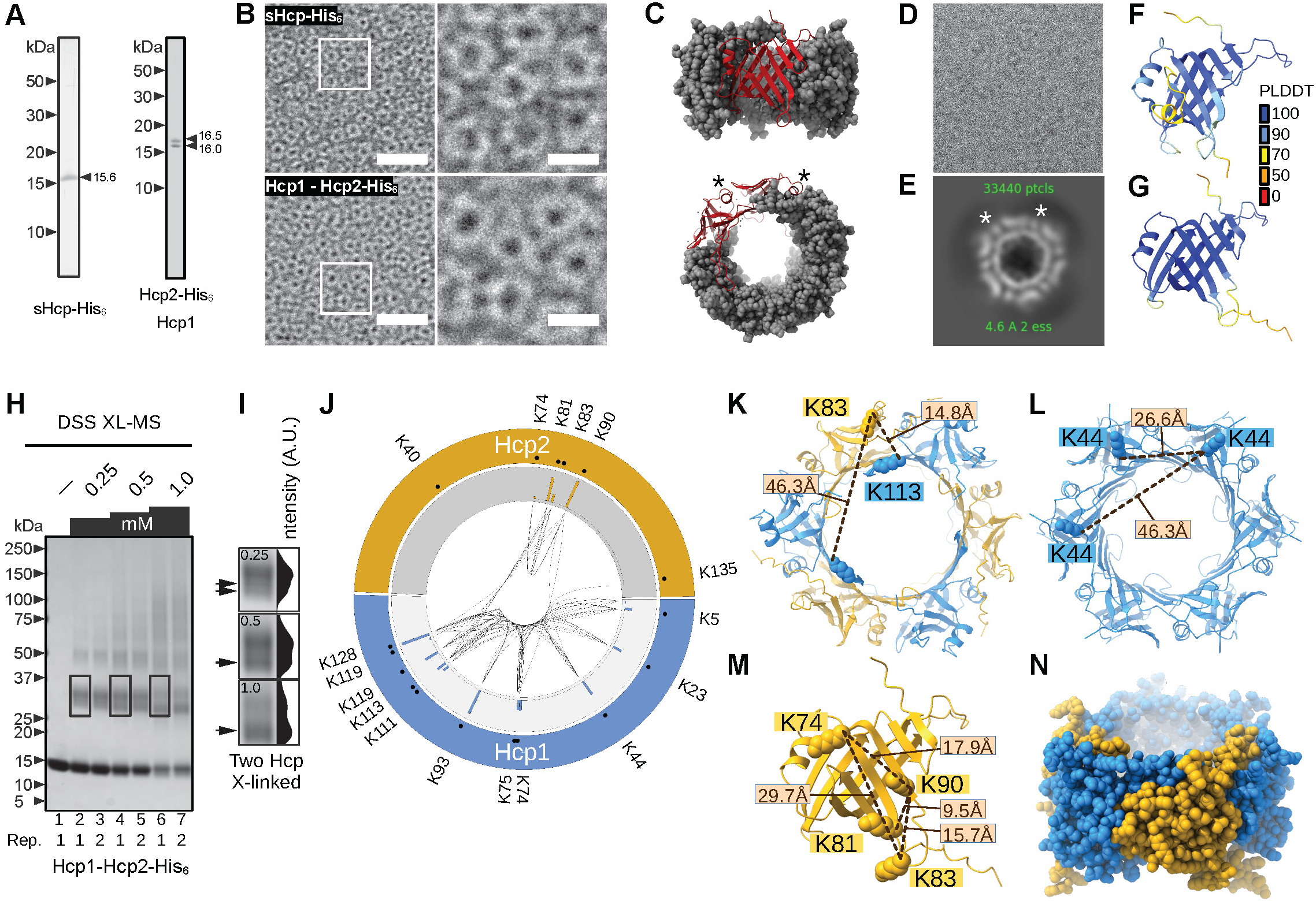
Purification and biochemical characterization of the major and minor Hcp subunits. **A)** Phosphoimager scan of SYPRO Ruby stained SDS-PAGE gels of purified sHcp-His6 and Hcp1 and Hcp2-His6 complexes. **B)** Negative staining electron microscopy image of sHcp-His6 and Hcp1 and Hcp2-His6 complex. Scale bars: 50 nm (left) and 15 nm (right). **C)** Crystall structure of sHcp. Asterisks denote two alpha helices in the crystal structure observed also by Cryo-EM data averaging. D) Cryo-EM data by single particle analysis of Hcp1-Hcp2-His6. **E)** Main class of particles isolated after a two-dimensional image single particle analysis). F) PLDDT colour-coded AlphaFold model of Hcp1, AF-Q5LDT5-F1. **G)** PLDDT colour-coded AlphaFold model of Hcp2, AF-A0A380Z0E1-F1. **H)** SDS-PAGE of two replicates (1 or 2) of X-linked Hcp proteins using DSS. **I)** Inset of SDS-PAGE with intensity line profile showing X-linked Hcp dimers at three DSS concentrations. **J)** Circos plot of all X-linked peptides found. Gray track contains a histogram of the total number of peptides found for each residue. **K)** Network of close residues in Hcp2 showing possible intramolecular X-links. **L)** K44 inter-protein Hcp1 X-links between residues at the same position modelled with AlphaFold as a Hcp1 homo-hexamer. **M)** Residues found in X-linked peptides between Hcp1 and Hcp2 modelled with AlphaFold as a 3:3 complex. **N)** AlphaFold model of an Hcp1-Hcp2 complex in a 4:2 ratio.

### XL-MS shows cooperatively formed adjacent Hcp1 protomers

To determine the spatial arrangement of individual protomers in the Hcp1‒Hcp2 complex, we performed cross-linking mass spectrometry (XL-MS). Copurified Hcp1-Hcp2-His^6^ was subjected to crosslinking with DSS, DSG or EGS at three concentrations (Figure 2H and Tables S2-S4). The crosslinked samples were observed via SDS‒PAGE, proteolyzed and subsequently analysed via mass spectrometry. SDS‒PAGE analysis of the control sample and crosslinked samples revealed the presence of a band migrating at the 15 kDa marker, corresponding to Hcp1 or Hcp2-His^6^ with either noncrosslinked subunits or with intraprotein crosslinks (Figures 2H lanes 1-3, 2I and S5). In the DSS crosslinked samples, a prominent band migrated between 25 kDa and 37 kDa, corresponding to two crosslinked Hcp proteins (Figure 2H, marked areas). At a low DSS concentration (0.25 mM), the intensity profile of the band showed faster migrating shoulder peaks, which became the most prominent band at higher concentrations (Figure 2I, arrows). This suggested that at higher crosslinker concentrations, two Hcp proteins were crosslinked by multiple linkers, which caused the two proteins to decrease their Stokes radii and migrate faster. To a certain extent, the band migrating close to the 50 kDa marker had the same appearance and showed the same behaviour at higher concentrations of the crosslinker. Although it was less clearly visible, we suggest that three Hcp proteins could also be crosslinked by multiple crosslinkers. Similar observations were made in the analyses of samples treated with DSG (Figure S5 and Table S3). At the tested concentrations, EGS did not efficiently generate crosslinked peptides (Figure S5 and Table S4). Above the 50 kDa marker, there were more discrete bands in the crosslinked samples, which decreased in intensity towards the higher-MW markers. This suggested that more than three Hcp protomers were crosslinked but at lower apparent frequencies. Quantification of both intra- and interprotein crosslinked peptides revealed that residues K119 in Hcp1 and K90 in Hcp2 were the most crosslinked residues (Figure 2J, histogram), suggesting that these residues are exposed. Filtering by residues that appear in multiple replicates (Table S5), the Hcp2(K83)-Hcp1(K113) crosslink was the most consistent crosslink (Figure 2K) among the hetero-interprotein crosslinks. This result confirms the expected interaction between Hcp1 and Hcp2. Consistent with our hypothesis that Hcp1 could form homointeractions in the presence of Hcp2, based on cooperative two-hybrid assays, we found multiple peptides across replicates showing interprotein Hcp1 crosslinks occurring between two identical residues in two Hcp1 subunits, *e.g.,* Hcp1(K44)-Hcp1(K44) and Hcp1(K113)-Hcp1(K113) (Figures 2L, S5 and Table S4). Peptides between residues K111 or K128 crosslinked to residues at the same position also occurred (Table S4). Interestingly, we were unable to observe any Hcp2 interprotein crosslinks, even though there was a network of exposed residues that consistently formed intraprotein crosslinks, namely, K74, K81, K83 and K90 (Figure 2M). In particular, Hcp2(K90) is very exposed, as shown by the total number of crosslinks, but Hcp2(K90) was never found in crosslinks with another Hcp2(K90). This result suggests that Hcp2 is not found close to another Hcp2 molecule or at least less frequently compared withs Hcp1 being close to another Hcp1.

To gain insight into the possible arrangements of the Hcp1-Hcp2 complex, we obtained hexameric models via the AlphaFold server (Figure S6). Using a Cα-Cα cut-off distance of 30 Å ^29^, we classified the observed crosslinks as permissive or non-permisisve in the models. We observed that in an Hcp1 homohexamer model, interprotein residues between K44 and K113 fell within the 30 Å distance in adjacent protomers (Figure 2L, S5 and Table S6) but not between residues two protomers away (P+2). Only the distance between Hcp1 K111 residues in protomer P and P+1 or P+2 was permissive for crosslinking. Conversely, in a Hcp1-Hcp2 (3:3) model in which Hcp1 protomers are two protomers away, only K111 was in the permissive distance. These results suggest that a structural model in which Hcp1 protomers are more frequently found adjacent to each other is more compatible with our crosslinking data. Interprotein crosslinking between Hcp1 and Hcp2, such as Hcp1(K113)-Hcp2(K83) (Figure 2K and Table S6), occurred only within the permissive distance when Hcp1 and Hcp2 were adjacent, confirming that neighbouring Hcp1‒Hcp2 proteins interact within the hexameric assembly. The distance between the observed Hcp2 crosslinked residues, with the exception of Hcp2(K74)-Hcp2(K90), was permissive only within the same protomer and not between adjacent protomers (Figure 2M and Table S6). This finding suggests that the majority, if not all, of the crosslinks observed between Hcp2 lysine residues were intraprotein. We measured the theoretical distance between all the residues found in crosslinked peptides and the same residue in the adjacent protomer in an Hcp2 homohexamer model (Table S6), and these residues were not permissive to crosslinking. Therefore, we cannot confirm that Hcp2 directly interacts with itself, as observed by two-hybrid assays, because even if Hcp2 assembles adjacent to another Hcp2, it could not be crosslinked at the same residue. However, the lack of interprotein cross-links in Hcp2 suggests that Hcp1 interprotein cross-links occurred among adjacent protomers and not randomly between Hcp protomers in two different hexamers. Interestingly, Hcp2(K40) did not show any crosslinks and indeed appeared buried in all the AlphaFold models when Hcp2 was adjacent to Hcp1 or Hcp2. K40 had a change in accessible surface area of -66.7 Å^2^ upon hexamer assembly, whereas K90 resulted in a change of only 0.4 Å2, indicating that K40 but not K90 is buried upon hexamerization (Figure S5). We were not able to explain two cross-linked residues involving Hcp1, K128 residues modeled at non allowed distances (Figure S5 and Table S6). However, the distance was close enough to be permissible, and these residues could be considered within range of other residues previously reported ^29^. Finally, data obtained from the published proteome of *B. fragilis* ^30^ revealed that Hcp1 is ∼53x more abundant than Hcp2 is (Figure S6), suggesting that Hcp2 is limiting. Notably, sHcp is ∼1000 fold more abundant than Hcp2 is, which is consistent with an inner tube structural role. Based on these differences in abundance, we refer to sHcp as the majour subunit and Hcp-1-4 as minor subunits. All these results, are consistent with an arrangement of the Hcp1‒Hcp2 hexamer, in which Hcp1 molecules are adjacent at higher frequencies and Hcp2 can be found in at least one or two copies, considering that the interaction between Hcp1 and Hcp1 depends on the presence of Hcp2, as shown by two-hybrid assays, and that Hcp1 and Hcp2 do not appear to be equimolar inside the bacterial cell. Therefore, we obtained a Hcp1-Hcp2 structure from the AlphaFold server with a 4:2 stoichiometry (Figure 2N and S6). The model showed an asymmetric structure with three adjacent Hcp1 protomers and the two Hcp2 protomers separated by another Hcp1. Notably, the Hcp2 protomers were not adjacently assembled. Analysing non-crosslinked samples (Table S7), showed that there were more Hcp1 peptides found compared to Hcp2. All these results are consistent with an Hcp1-Hcp2 complex model formed primordially of Hcp1 with fewer Hcp2 subunits.

### The Hcp1‒Hcp2 complex forms heterohexamers of variable stoichiometry

Having stated that Hcp1-Hcp2 forms ring-shaped hexamers, we set out to determine the stoichiometry of this complex. To this end, we repeated the purification of Hcp1-Hcp2-His^6^, this time eluted with an imidazole step gradient and analysed the fractions via SDS‒PAGE densitometric analysis. Three fractions were collected at each imidazole concentration. As a control, we constructed a plasmid expressing sHcp-sHcp-His^6^, which should promote the formation of mixed hexamers of tagged and untagged monomers. Indeed, SDS‒PAGE analysis of the eluted sHcp-sHcp-His^6^ bands revealed two fast-migrating bands corresponding to the tagged and untagged proteins (Figure 3A). Two slower migrating bands appeared just below the 50 kDa marker with densities proportional to the corresponding higher-migrating bands, which we interpret to be SDS-resistant trimers of sHcp (labelled sHcp*) because single bands appeared in analysis of purified sHcp-His6 samples just above the 15 kDa and just below the 50 kDa markers (Figure S4D). Higher proportions of sHcp-His^6^ eluted with increasing concentrations of imidazole, possibly because hexamers containing more His^6^-tagged protomers bind to the Ni-NTA resin with more avidity and required more imidazole to elute (Figure 3A). Different results were obtained for the Hcp1-Hcp2-His^6^ complex (Figure 3B). First, no SDS-resistant proteins were eluted. Second, most of the protein peak was eluted at 150 mM imidazole rather than at 250 mM imidazole as for the sHcp-sHcp-His6 complexes. Given that homohexameric sHcp complexes could contain up to six His-tagged subunits, whilst in the case of untagged Hcp1 with tagged Hcp2 six His-tagged subunits (and thereby the same binding strength to the resin) would be observed only if hexamers devoid of Hcp1 formed. Therefore, lower imidazole elution suggested that Hcp1-Hcp2-His^6^ complexes did not have as many His^6^-tagged protomers as sHcp-sHcp-His^6^, which is consistent with the formation of Hcp1-Hcp2 heterohexamers. Third, at all elution concentrations, untagged Hcp1 appeared to be present in higher proportions than Hcp2-His^6^.

**Figure 3.**
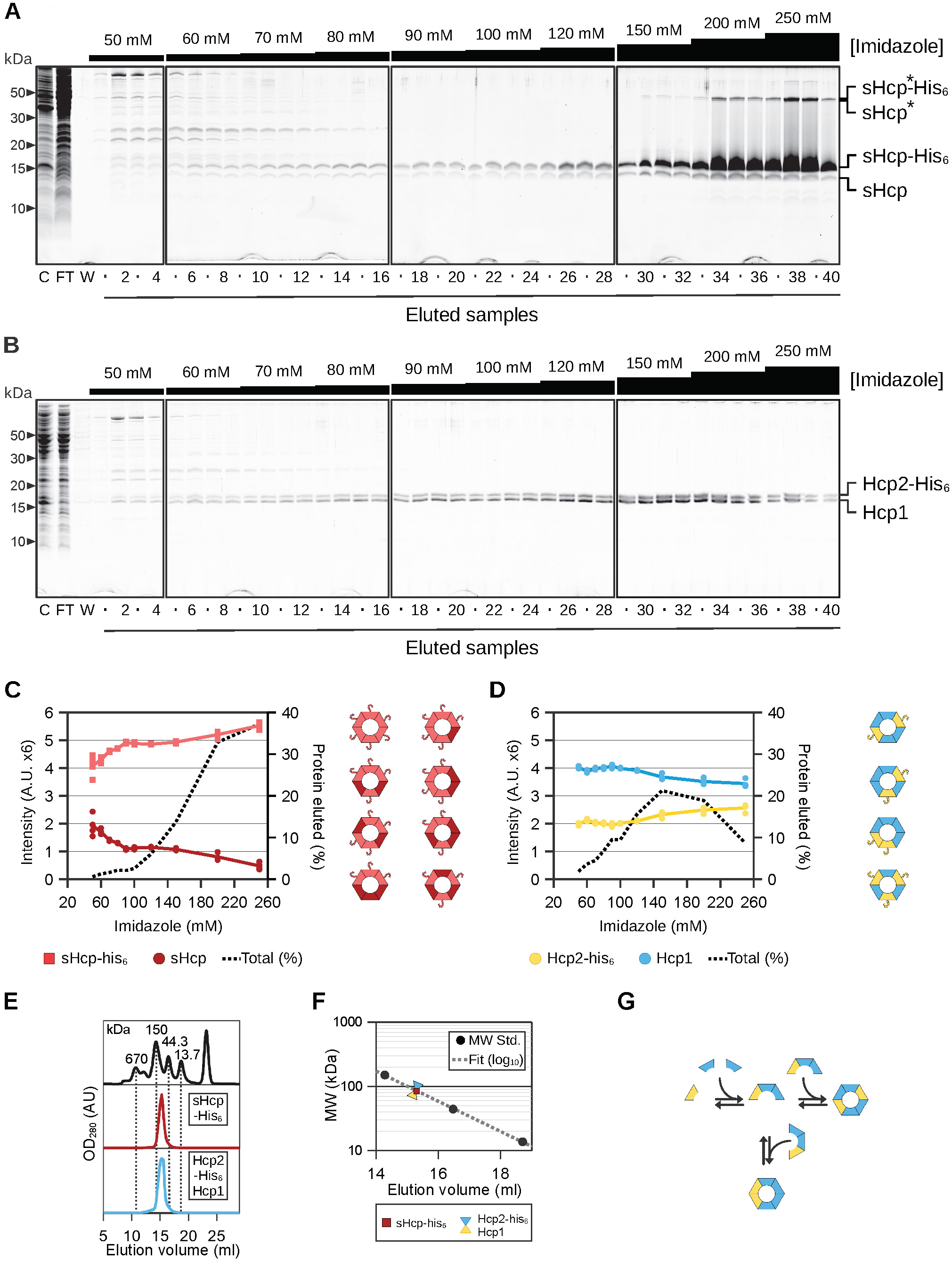
Stoichiometry of purified Hcp complexes as determined by an imidazole step gradient. A-B) SYPRO Ruby stained SDS-PAGE gels of samples obtained from the affinity co-purification of sHcp and sHcp-His6 or Hcp1 and Hcp2-His6. Four elution steps were performed for each imidazole concentration (black rectangles on top of the gels). Labels: C (clear fraction), FT (flow-trough), W (wash), 1-40 (elution fractions). **C-D)** Densitometric quantification of SYPRO Ruby fluorescence intensity from the gels shown in A and B. The average fractional density of fluorescence intensity of the two regions of interest (ROI) corresponding to each purified Hcp protein (with and without His6 tag) normalized to the total density multiplied by 6 (x6) is plotted against the imidazole concentration. The percent of eluted protein is shown in the secondary *y*-axis as the sum of the two ROIs in each lane, divided by the sum of all ROIs and multiplied by 100. The possible stoichiometric arrangements of sHcp-sHcp-His6 and Hcp1-Hcp2 complexes are indicated by color coded filed hexamers: Light red (sHcp-His6), dark red (sHcp), yellow (Hcp2-His6) and skyblue (Hcp1). **E)** SEC analysis of a protein-standard mixture (top) purified sHcp-His6 (midddle, red) and the co-purified complex of Hcp2-His6 with Hcp1 (bottom, blue). **F)** Calibration of size exclusion chromatography molecular weight standards. Elution volumes of the SEC molecular weight protein-standards (black circles) are plotted against their molecular weights. A linear fit using three of the log10 MW of the standards (dotted line) and the extrapolated MW of the Hcp complexes are also shown. The intrapolated MW of sHcp-His6 was 85.1 kDa (red square). The intrapolated MW for the Hcp2-His6 co-purified with Hcp1 was 86.8 kDa (yellow-blue triangles). **G)** Assembly of possible radially asymmetric heterohexamers of Hcp1 and Hcp2.

To obtain quantitative measurements of the stoichiometry of the Hcp complexes, we performed densitometric analysis of the SDS‒PAGE SYPRO-ruby-stained gels. We calculated the proportions of tagged and untagged proteins and normalized them to six (under the assumption that each eluted particle is a hexamer). For sHcp-sHcp-His^6^, only the fast-migrating bands were analysed, which revealed that the number of eluted sHcp-His^6^ steadily increased in proportion to the imidazole concentration (Figure 3C). However, for the Hcp1-Hcp2-His^6^ complex, up to 120 mM imidazole, the eluted complexes had a 4:2, Hcp1:Hcp2 ratio (Figure 3D). Above 120 mM, the complexes had fractional ratios, which we interpret to be mixtures of mostly 4:2 with some 3:3 complexes, although we cannot exclude other ratios or trace amounts of supracomplexes. Additionally, we quantified the total fraction of protein eluted at each imidazole concentration by adding the density of both the untagged and tagged proteins and dividing by the sum of all the densities (Figure 3C and 3D, dashed lines). This analysis revealed that approximately half of the eluted Hcp1-Hcp2-His^6^ had a 4:2 ratio and confirmed that the protein peak of elution was at 150 mM imidazole.

To obtain full insight into the stoichiometry of the Hcp1-Hcp2-His^6^ complex, we subjected it to analytical size exclusion chromatography (SEC) with sHcp-His^6^ as a control. Using molecular weight standards, we observed that sHcp-His^6^ eluted as a single peak from the column after 15 ml between the 150 and 44.3 kDa markers, similarly to the Hcp1-Hcp2-His^6^ complex (Figure 3E). To better quantify the size of the complex, we selected three markers from the molecular weight (MW) standards and plotted the MW *vs.* the elution volume, which showed a logarithmic relationship. The MW values of sHcp-His^6^ and Hcp1-Hcp2-His^6^ extrapolated from the plot were 85.1 and 86.8 kDa, respectively (Figure 3F). The calculated MWs from the sequence for one sHcp-His^6^ hexamer are 93.6 kDa, while 97 kDa for a 4:2 Hcp1-Hcp2-His^6^ complex. This shows that the Hcp1-Hcp2-His^6^ complex is formed by hexamers. Together with the findings described above, these results suggest that the Hcp1-Hcp2-His^6^ complex preferentially assembles *in vitro* into hexamers with a stoichiometry of 4:2. This implies that hexamers of Hcp1-Hcp2 complexes must contain adjacent Hcp1 protomers, which were observed in two-hybrid assays, but only if Hcp2 was present. Overall, we propose that the Hcp1-Hcp2 complex may form through cooperative interactions via Hcp1 homointeractions mediated by Hcp2 (Figure 3G). Moreover, we conclude that *in vitro* assembled Hcp1‒Hcp2 complexes have variable stoichiometries.

### Minor Hcps are linked to effector recognition and secretion

Our data show that the product of *hcp1* forms chimeric complexes with the product of *hcp2*, which is the gene located downstream. Interestingly, the effector *bte1* is downstream of *hcp2*. Bte1 was recently described as the first T6SS effector with the capacity to kill bacteria by altering protein folding ^31^. As reported by Coyne *et al.* ^16^, in the high-synteny arrangement of *Bacteroidales* GA3, two effectors are encoded in variable cassettes but are always arranged in the same order. In most strains one effector is downstream of an Hcp2 homologue. Therefore, we hypothesised that the Hcp1-2 complex could interact with Bte1. First, we tested the interactions of Bte1 with all minor Hcp proteins via bacterial two-hybrid assays. Our results revealed that Bte1 fused to T18 or T25 fragments interacted with both Hcp1 and Hcp2 fused to T18 or T25 fragments when Hcp2 and Hcp1 were coexpressed, but not with the Hcp3 or Hcp4 T18 and T25 fusions (Figure 4A). To confirm this interaction, we copurified a His-tagged version of Bte1 with Hcp1 and Hcp2 and analysed it via size exclusion chromatography. Two major peaks were observed (Figure 4B). Pooled fractions from each peak were subsequently analysed by SDS‒PAGE, which revealed that peak 1 corresponded to the Hcp1‒Hcp2‒Bte1 ternary complex, whereas peak 2 corresponded to Bte1-His^6^ alone (Figure 4C). Negative staining electron microscopy images of the ternary complex showed particles in various orientations (Figure 4D). A few particles were filled with a density in the centre (Figure 4D, asterisks), and other particles appeared as empty rings (Figure 4D, empty circles). These results suggest that the minor Hcp heterohexamers can act as recognition particles in the case of *Bacteroidales* T6SS effectors.

**Figure 4.**
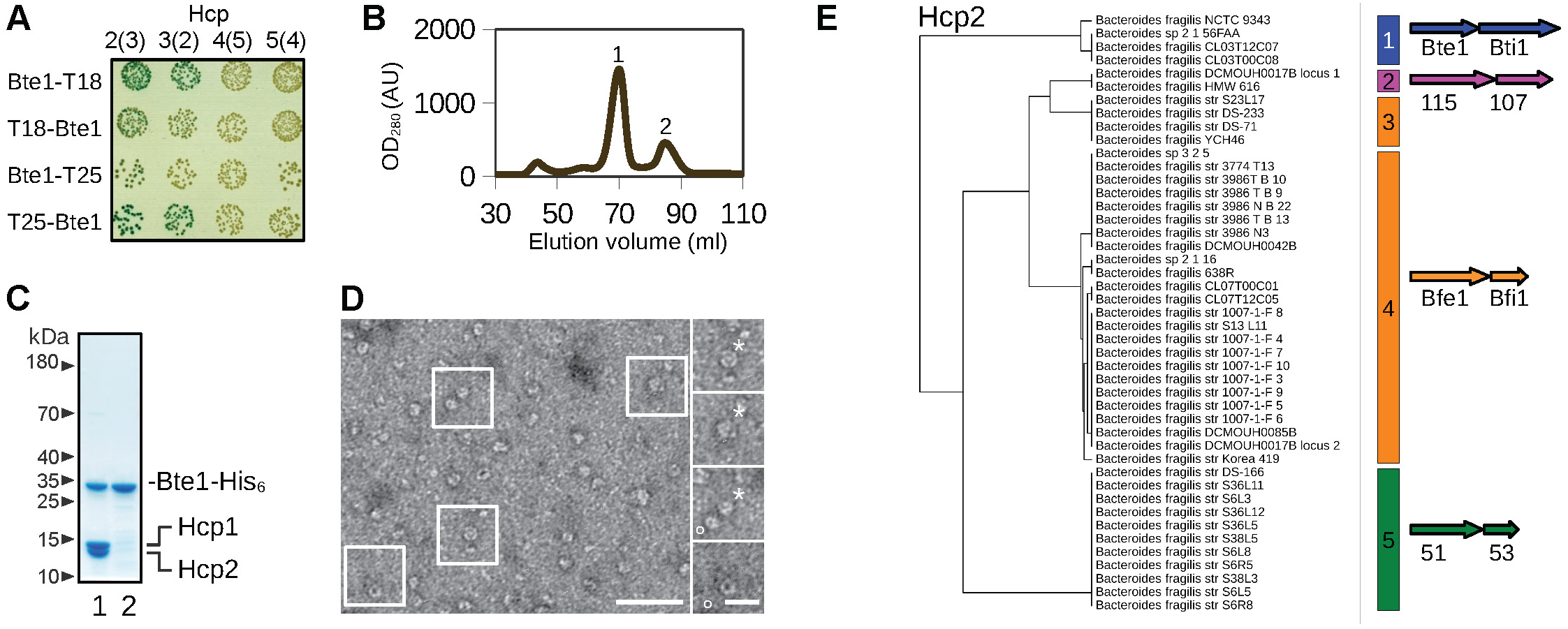
Hcp2 determines binding to GA3 T6SS effectors. **A)** Photographic images of cooperative two-hybrid assay for pairwise interactions between T18 or T25 translational fusions of Bte1 with Hcp1 in the presence of Hcp2 (2(3)), Hcp2 in the presence of Hcp1 (3(2)), and so on. Pictures were taken at 36h. **B-C)** Size exclusion chromatography of co-purified Hcp-effector complex. Two peaks were characterized by SDS-PAGE. The first and largest peak (1) contained Hcp1 (16.0kDa), Hcp2 (15.3 kDa) and Bte1-His6. The second peak (2) only contained the effector. **D)** Negative staining electron microscopy of the Hcp1-3-Bte1 effector complex. Filled structures are marked with asterisks (*), empty structures are marked with empty circles (○). Scale bar 50 nm (insets 20 nm). **E)** Clustering of phylogenetic tree of Hcp2 homologues from GA3 T6SS with their cognate effector immunity pairs. Bte1 with Bti1 (blue), cluster 115 with immunity gene (magenta), Bfe1 with Bfi1 (orange) and cluster 51 with immunity gene (green).

Three more effectors were described for the T6SS GA3 in other strains in addition to Bte1 encoded in the minor *hcp*-adjacent variable region. Two proteins were initially annotated using Hidden Markov models (HMMs) and categorized as cluster 51 and cluster 115 (using Coyne *et al.* clustering nomenclature ^16^), which were upstream of immunity proteins named clusters 107 and 53, respectively. The third type of effector, present in *B. fragilis* 638R, is called *bfe1* ^20^ and has been shown to target other *Bacteroides* spp. To gain an overview of the probability that the *Bacteroidales* minor Hcp proteins function as effector recognition particles in general, we generated a phylogenetic tree of all GA3 available *Hcp2* genes described in Coyne *et.al* ^16^. The Hcp2 homologues had a high degree of identity, but they were separated into five clusters (Figure 4E and S8A). We then analysed the effectors adjacent to each cluster and observed that the four types of effectors segregated in different clusters (Figure 4E). Only Bfe1-like effectors, similar to the effector found in *B. fragilis* 638R, which were the most common, segregated in two clusters, which were closer to each other. This cosegregation did not occur when sHcp was used to generate the phylogenetic tree, suggesting that sHcp is not a recognition factor for T6SS GA3 effectors (Figure S7A). Hcp1 and Hcp3 also cosegregated with their effector–immunity pairs (Figure S7B-C). The clustering of Hcp2 with effectors was also present in T6SSs with GA2 clusters (Figure S8B) and segregation of Hcp sequences with effectors was described by Bao *et al.* ^23^ Considering that Bte1, Hcp1 and Hcp2 were found to be secreted ^19^, it is reasonable to propose that they are secreted as a complex. Taken together, our results are consistent with the idea that minor Hcp proteins are cognate to effectors and participate in secretion, and that this feature is conserved in *Bacteroidales* T6SSs.

## DISCUSSION

This study revealed that in Bacteroidales T6SSs there are different types of Hcp subunits with specific assembly properties and functions. Firstly, a role for sHcp as major subunit constituting the bulk of the inner tube is consistent with data showing its high abundance and its capacity to form homohexamers. In the case of Hcp1 and Hcp2, we showed here that these minor subunits form heterohexamers and have a specialized function of recognition particles for effector secretion.

Our results are consistent with and extend the findings of Silverman *et al.* and Howard *et al.*, who delineated a role for Hcp proteins in effector recognition ^13,14^. In the case of *Pseudomonas aeruginosa,* Hcp subunits have been shown to recognize effectors as chaperones but are also required for export ^13^, and different T6SSs present in this bacterium can specifically recognize cognate effectors ^14^. However, our results suggest a clear division of functions between major and minor Hcp subunits. Considering that effectors in gut Bacteroidales are encoded in variable chromosomal regions within T6SS loci, our results imply that the recombination of effector genes require co-segregation of genes encoding cognate Hcp proteins that would facilitate export of multiple effectors by the same secretion system, suggesting a bypass of the evolution of specific recognition motifs only to major Hcp, VgrG or PAAR subunits. Furthermore, our results imply that the functional characterization of many Bacteroidales T6SS effectors may require the characterization of their cognate Hcp protein complexes.

Consistent with a structural role for sHcp, proteomic data showed that this subunit is ∼1000-fold more abundant compared to Hcp2 ^30^, which is compatible with this protein constituting the bulk of the inner tube and that Hcp1 and Hcp2 act as minor Hcps where Hcp2 is an effector-specific limiting factor. These data are consistent with previously published RNA-seq data ^20^ showing that *shcp*’s mRNA is up to ∼43 fold more abundant than the minor *hcp’s* mRNAs, suggesting that rate of transcription and/or mRNA lifetime modulation may contribute to inner tube subunit abundance. Moreover, sHcp is the only subunit present in all three genetic architectures of Bacteroidales T6SSs, whereas only GA2 and GA3 have multiple minor Hcp subunits ^16^. T6SSs with GA1 have two copies of sHcp with no adjacent genes coding for effectors ^16,23^, suggesting that effector recognition and recombination may function differently in organisms with these T6SSs.

Two-hybrid assays showed a high level of interaction among Hcp subunits. However, in competition assays Hcp1 preferentially bound to Hcp2 and Hcp3 preferentially bound to Hcp4. These interactions coincide with the corresponding two pairs of genes being adjacent and are consistent with a model in which these proteins coevolve to form heterohexamers that recognize other proteins. This finding is further supported by a high level of synteny in gut Bacteroidales T6SS loci. We also identified other interactions that await biochemical confirmation. For example, sHcp could bind to Hcp1 or Hcp3. These minor Hcps may intersperse with major subunits in the assembled inner tube. However, the difference in abundance and the greater tendency of sHcp to form homohexamers may increase the difficulty of detecting these interactions both *in vivo* and *in vitro*. We also observed binding between Hcp1 and Hcp4. Hcp4 was not detected in proteomic data ^30^, suggesting that, like Hcp2, Hcp4 may be present in low abundance. It could be that Hcp1 may carry both Hcp2 and Hcp4 protomers to mediate specific interactions. Furthermore, it is feasible that these complexes have additional interaction partners *in vivo.* How the minor Hcp subunits assemble within the inner tube and bind to VgrG remains to be elucidated, but the proteomic abundance data suggest that the formation of the Hcp1‒Hcp2 complex and effector recognition is limiting. Therefore, there may be a specific assembly pathway for minor Hcp subunits and their binding partners to assemble at the tip of the inner tube that ensures their function in secreting cognate effectors.

Copurification, electron microscopy imaging and SEC of the Hcp1-Hcp2 complex demonstrated that it formed heterohexamers. This type of assembly has implications for T6SS containing specialized or evolved Hcp subunits with effector-carrying C-terminal extensions ^24,25^, which are also present in gut *Bacteroides* ^16^. Our results imply that in systems carrying evolved Hcps, major and minor Hcps carry distinct functions forming distinct assemblies for secretion.

Our results also highlight the question of how hexameric Hcp complexes are assembled. The SDS-resistant Hcp trimers observed during sHcp purification could represent a stable intermediate in its assembly pathway. Therefore, our results hint that two stable trimer intermediate structures assemble into hexamers. The heteromeric assembly of the Hcp1-Hcp2 complexes suggests a mechanism by which effectors are concomitantly assembled with minor Hcps. Indeed, the homointeractions between Hcp1 and its purification required the presence of Hcp2, and it is reasonable to think that Bte1 recognition may benefit from such a cooperative assembly pathway. He *et al.* discovered a conformational change associated with sHcp binding to VgrG ^28^. The question of how the Hcp1-Hcp2 complex is assembled in the context of the whole secretion system remains open.

In polymorphic toxins, the N-termini typically carry secretion specificity domains and the C-termini carry toxin domains. Toxins are frequently encoded in gene fragments located adjacently upstream of genes encoding cognate antitoxin proteins ^32^. Such synteny has resulted in toxin-antitoxin (immunity) pairs being acquired by horizontal gene transfer as gene fusions of N-terminal secretion specificity domains ^33^. The high level of synteny of the *Bacteroides* T6SSs translates into function by linking the position of the gene and the hierarchy of interaction of their products as shown here by protein-protein interactions between Hcp1, Hcp2 and Bte1 (which in turn would interact with its antitoxin, Bti1). Therefore, our results suggest that in the case of *Bacteroidales* T6SSs, the minor *hcp* genes partly encode domains that mirror transcriptional fusions of polymorphic toxins encoding secretion specificity domains, although without being fused. Indeed, Hcp2 homologues clustered according to their cognate effectors in the phylogenetic tree, suggesting that Hcp2 is a cosegregated specificity factor. In a previous Hcp study, Howard *et al.* concluded that the interactions between effectors and cognate Hcp proteins depend on the overall Hcp structure and not on specific residues on the basis of the clustering of specialized and core Hcps performed on phylogenetic analyses of a large subset of proteins ^14^. However, our phylogenetic analysis using a specific subset of *Bacteroidales* T6SS GA3 Hcps clearly segregated effectors with cognate minor Hcps but not with major sHcp proteins. This model is consistent with the minor Hcps participating in specific recognition of effectors. If this is true, it could mean that minor Hcps can recognize different effectors with a minimum of variability, encoded in very few residues. Although this specificity could be also encoded through selective steric hindrance as suggested by Howard *et al.* ^14^. This sort of specificity could be aided by an asymmetric assembly with limiting Hcp2 subunits as recognition factors, which could accommodate effectors in the minor Hcp complex with specific stoichiometry.

It was recently reported that Bte1 is an effector that disrupts protein folding in the periplasm and is used for antagonistic interactions ^31^. Lim *et al.* showed that heterologous expression of Bte1 in *Bacteroides thetaiotaomicron* causes toxicity by targeting the chaperone-acting PpiD-YfgM complex, thus suggesting that Bte1 requires no other factors for its activity. Moreover, Hcp1, Hcp2 and Bte1 have been shown to be present in the secreted fraction ^19,21^. Therefore, it is possible that Bte1 is co-translocated with Hcp1 and Hcp2 as a complex. Alternatively, a chaperone function could be necessary to position Bte1 for secretion.

The separation of major and minor subunit proteins with high and low abundance, respectively, in gut Bacteroidales T6SSs appears akin to macromolecular export systems that assemble filamentous structures. For example, type II secretion systems include one major pseudopilin (or endopilin) and four minor pseudopilins. The minor subunits form a tip complex that functions in priming the assembly of the major pseudopilin into a filament ^34,35^. This type of assembly and function of minor pilins is conserved in type IV pili ^36–38^. In type 1 fimbria minor pilins mediate adhesion and appear at the tip of fimbriae because of their higher affinity to the usher subunit ^39,40^. Further studies are needed to elucidate the *in vivo* assemblies of minor Hcp subunits in the *Bacteroides* T6SSs. Finally, our results shed light on how Bacteroidales T6SSs evolved to exhibit a high diversity and variability of effectors. The presence of major and specialized minor Hcp subunits that recognize these effectors is an evolved feature that facilitates this variability, contributing to *Bacteroides* biodiversity.

## MATERIALS AND METHODS

### Bacterial two-hybrid assay

Twenty-five microlitres of ultracompetent *E. coli* BTH101 (*cyaA*^-^) cells were cotransformed with plasmids carrying T18 and T25 translational fusions via heat shock. After recovery in 2X YT medium, 10 microlitres of the bacterial suspension were spotted on Luria broth agar plates containing 40 μg/ml and 0.125, 0.25 or 0.5 mM IPTG. Drops of bacterial suspensions cotransformed with control plasmids were included in each plate. The plates were incubated at 30 or 37 °C for 24 and 36 h. For image analysis (see below), the density of bacteria must be similar. We found that the best results were obtained when the drops of bacteria grew between single colonies and a small lawn. The plates were photographed at each time point via a Fas-DIGI system (NIPPON genetics) equipped with a white screen and a PENTAX RICOH-IMAGING camera with the following settings: ISO 200, F number 2.5, and an exposure time of 1/800 s. The original images look dim, but this dimness is important for image analysis. The images were analysed with FIJI by separating the colour image into its three RGB components and taking the red channel only. The image was inverted, and the average value of the background outside the plates was manually subtracted. To quantify each spot, a region of interest (ROI) that was only slightly smaller than the growing bacteria excluding the borders but equal for each spot was measured for each sample via Fiji to obtain the average intensity. The normalized intensity at each spot on the plate was calculated by subtracting the value of the negative control from the intensity value of each spot and dividing it by the difference between the positive and negative controls. For some experiments, two pictures were taken for each plate. The two pictures were treated as above before quantification and placed in a stack in FIJI. The two images were aligned via the linear stack alignment (SIFT) plugin and stack-averaged. This process greatly reduced differences in the background caused by deviations in light capture at certain pixel position in the camera’s sensor, which caused some spots to appear to have less signal than the negative controls.

### Protein purification

Protein purification was carried out by expressing *hcp* genes from pET28 constructs in BL21(DE3) cells. Co-expression of *Hcp1, Hcp2* and *bte1* was performed using plasmid pETDUET-1. Cells were grown in LB medium at 37°C until exponential phase and induced with 0.4 mM IPTG after cooling the cells. Induction was performed overnight at 20°C or for three hours at 30°C. Cells were harvested by centrifugation and resuspended in 50mM NaP 8.0. 500mM NaCl, 10% glycerol, 10mM Imidazole, 1mM β-mercaptoethanol. Lysis was performed by sonication and the cleared lysate was prepared by centrifugation at 40,000 g for 30 minutes. The cleared lysate was incubated with a Ni-NTA resin (ThermoScientific) for 1 h, washed with resuspension buffer and eluted with resuspension buffer containing 300 mM imidazole. For preparative size exclusion chromatography (SEC) a HiLoad 16/600 Superdex200pg column (Cytiva) was used. For analytical SEC, a Superdex200 10/300gl column (Cytiva) was used and a Protein Standard Mix 15 - 600 kDa (Merck). Proteins samples were characterised using SDS-PAGE stained with blue Coomassie or SYPRO-Ruby.

### Densitometric analysis of the SDS‒PAGE gel (fluorescence intensity)

The gels were stained with SYPRO-Ruby and scanned with a GE Typhoon FLA 9000 Gel Imager. The tiff images were then analysed in FIJI. For each quantified protein band, two ROIs were created. The first ROI contained the band. The second ROI was the same size but encompassed an immediately adjacent area without stain as a blank. The ‘integrated density’ of each ROI was measured, and the blank was subtracted to obtain the corrected fluorescence intensity (fluo^corr^) of the protein band. To obtain the normalized SYPRO-ruby fluorescence intensity at each lane (fluo^norm^) of each band of His-tagged (fluo^norm^1) protein *vs.* non-His-tagged protein (fluo^norm^2), the corrected fluorescence intensity of each band was normalized by the following equation: fluo^norm^1= fluo^corr^1/(fluo^corr^1 + fluo^corr^2)*6 or fluo^norm^2= fluo^corr^2/(fluo^corr^1 + fluo^corr^2)*6. The obtained number represents the fraction multiplied by six of one or the other protein normalized by the total intensity.

### Electron microscopy

For transmission electron microscopy (TEM) the samples were resuspended in purification buffer and 3.5 μL adsorbed to formvar and carbon-coated glow discharged grids (cupper, 300 mesh, Ted Pella Inc.) and were negatively stained with 1.5% w/v uranyl acetate. The samples were imaged with a Talos L120 TEM using the Ceta detector and Velox software (Thermo Fisher). For Cryogenic TEM (Cryo-EM) 4 μL of the samples were adsorbed to holey carbon film on 300 mech cupper grids (Quantifoil). The Grids were blotted and plunge frozen in liquid ethane using the Vitrobot (Thermo Fisher). Cryo-EM imaging was performed with Titan Krios G2 Cryo-EM using Gatan K2 BioQuantom detector and EPU software (Thermo Fisher). The Cryo-EM images were analysed by single particle analyse approaches with the CryoSPARC software package.

### Protein alignments and phylogenetic analyses

Protein alignments were performed at the MAFFT website ^41^ using the G-INS-I algorithm and UPGMA. The output phylogenetic guide trees with effector or Hcp alignments were plotted with TreeViewer ^42^. Effectors were clustered by UPGMA distance in JalView.

### Crosslinking of the Hcp1-Hcp2 complexes

A total of 20 μG of Hcp1-Hcp2 complexes in 20 mM NaP, pH 8.0, 150 mM NaCl, 1 mM DTT were crosslinked in duplicates with three different concentrations (0.25 mM, 0.5 mM and 1.0 mM) of heavy/light DSS (DSS-H12/D12, Creative Molecules Inc., 001S), heavy/light DSG (DSG-H6/D6, Creative Molecules Inc., 010S) or heavy/light EGS (EGS-H12/D12, Creative Molecules Inc., 003S), respectively in a total reaction volume 20 uL. Non-crosslinked samples were kept as controls. All samples were incubated for 30 min at 37°C, 1000 rpm. The cross-linking reaction was quenched with a final concentration of 50 mM of ammonium bicarbonate for 15 min at 37°C, 1000 rpm. For SDS-PAGE analysis of the crosslinked samples, 5 uL of the above reactions were mixed with 5 uL of 2x sample buffer (BioRAD) and boiled at 95°C, 5 min. The samples were analyzed on a 4-20% BioRAD Criterion TGX gel.

### Sample preparation for mass spectrometry

The remaining samples crosslinked with DSS and DSG were denatured using an 8 M urea– 100 mM ammonium bicarbonate solution. For the EGS-crosslinked samples, 8 M urea without ammonium bicarbonate was used. The cysteine bonds were reduced with a final concentration of 5 mM Tris (2-carboxyethyl) phosphine hydrochloride (TCEP, Sigma, 646547) for 60 min at 37°C, 1000 rpm and subsequently alkylated using a final concentration 10 mM 2-iodoacetamide for 30 min at 22°C in the dark. For digestion, 1 μG of lysyl endopeptidase (LysC, Wako Chemicals, 125-05061) was added, and the samples incubated for 2 h at 37°C, 100 rpm. The samples crosslinked with DSS and DSG were diluted with 100 mM ammonium bicarbonate to a final urea concentration of 1.5 M, whereas the samples crosslinked with EGS were diluted using LC-H^2^O. For digestion, 1 μG of sequencing grade trypsin (Promega, V5111) was added for 16 h at 37°C, 1000 rpm. The digested samples were acidified with 10% formic acid to a final pH of 3.0. Peptides were purified and desalted using C18 reverse phase columns (The Nest Group, Inc.) following the manufacturer’s recommendations. Dried peptides were reconstituted in 50 μL of 2% acetonitrile and 0.1% formic acid prior to MS analysis.

### Liquid chromatography tandem mass spectrometry (LC-MS/MS)

A total of 1 μL of peptides were analyzed on an Orbitrap Eclipse mass spectrometer connected to an ultra-high performance liquid chromatography Dionex Ultra300 system (both Thermo Scientific). The peptides were loaded and concentrated on an Acclaim PepMap 100 C18 precolumn (75 μm × 2 cm,) and then separated on an Acclaim PepMap RSLC column (75 μm × 25 cm, nanoViper,C18, 2 μm, 100 Å) (both columns Thermo Scientific), at a column temperature of 45°C and a maximum pressure of 900 bar. A linear gradient of 2% to 25% of 80% acetonitrile in aqueous 0.1% formic acid was run for 100 min followed by a linear gradient of 25% to 40% of 80% acetonitrile in aqueous 0.1% formic acid for 20 min. One full MS scan (resolution 120,000; mass range of 400-1600 *m*/*z*) was followed by MS/MS scans (resolution 15,000) with a 3 sec cycle time. Precursors with a charge state of 3-8 were included. The precursor ions were isolated with 1.6 *m*/*z* isolation window and fragmented using higher-energy collisional-induced dissociation (HCD) at normalized collision energies (NCE) of 21, 26, 31. The dynamic exclusion was set to 45 s.

### Crosslinking data analysis

All spectra from the crosslinked samples were analyzed using pLink 2 (version 2.3.11). The target protein database contained the sequence for Hcp1 and Hcp2 and common contaminants. pLink2 was run using default settings for conventional HCD DSS-H12/D12, DSG-H8/D8 or EGS-H12/D12 crosslinking, with trypsin as the protease and up to 3 missed cleavages allowed. Peptides with a mass range of 600–6000 *m*/*z* were selected (peptide length 6–60 residues) and the precursor and fragment tolerance were set to 20 and 20 ppm, respectively. The results were filtered with a filter tolerance of 10 ppm and a 5% FDR. The mass spectrometry data has been deposited to the ProteomeXchange consortium via the MassIVE partner repository (https://massive.ucsd.edu/) with the dataset identifier PXD055833. The non-crosslinked control samples were analyzed in MaxQuant (v. 2.6.2.0) against the same sequence database as above. Fully tryptic digestion was used allowing two missed cleavages. Carbamidomethylation (C) was set to static and protein N-terminal acetylation and oxidation (M) to variable modifications. Mass tolerance for precursor ions was set to 10 ppm and for fragment ions to 0.02 Da. The protein false discovery rate was set to 1%.

### Circos plots and structural analysis

Circos plots were obtained in Rstudio using the Circlize library ^43^. The AlphaFold server ^44^ was used to create hexamer structures using default parameters and a random seed number. Visualization of the structures, distance measurements between residues and structural alignments were performed in Chimera ^45^.

## ACKNOWLEDGEMENTS

We acknowledge support from Mikael Lindberg and Lotta Happonen at the the Umeå node of the Protein Expertise Platform (PEP) /Protein Production Sweden (PPS) and the Swedish National Infrastructure for Biological Mass Spectrometry (BioMS)/ Integrated Structural Biology (ISB) platform and the SciLifeLab, respectively. We are thankful to Dr. Andrea Puhar, WWIEM, Queen’s University Belfast, UK for critical reading. Electron microscopy instruments and methods were supported by Michael Hall at the Umeå Centre for Electron Microscopy (UCEM)/ SciLifeLab national Cryo-EM infrastructure supported by the Kempe foundations and Knut and Alice Wallenberg foundations. B.E.U. received support from the Swedish Research Council (2019-01720, 2016-06598), Kempestiftelserna (JCK-1527.1, JCK-1724) and the Faculty of Medicine, Umeå University (Insamlingsstiftelsen grant 2021-2023). D.A.C. received support from the Kempestiftelserna (SMK-1860), Carl Tryggers Stiftelse för Vetenskaplig Forskning (CTS 15-96 and CTS 18-65) and the Faculty of Medicine, Umeå University (Insamlingsstiftelsen 2019-2021).

## CONTRIBUTIONS (CRediT taxonomy)

Conceptualization, data curation, formal analysis and writing original draft: DAC. Funding acquisition, supervision, resources and project administration: DAC & BEU. Visualization: SGSM, MKSAA, DAC. Methodology, Investigation, review and editing: SGSM, MKSAA, EJ, AK, US, PB, LS, BEU & DAC

**The authors declare no competing interests**

## Supplemental Information

### Supplemental methods

#### Crystallization of Se-Met labelled sHcp protein

The sHcp protein was genetically modified to contain an uncleavable C-terminal 6His-tag for immobilized metal affinity chromatography (IMAC) purification. For X-ray crystal structure determination, the sHcp protein was labelled with Seleno-Methionine (Se-Met) as described elsewhere^1,2^ for Single-wavelength Anomalous Diffraction (SAD) data collection.

Crystals of the C-terminal 6His-tagged Se-Met sHcp1 protein (amino acid sequence below) were set-up by mixing equal amounts, typically 2µl + 2µl of ultra-pure sHcp protein, at a concentration of 14 mg/ml, with precipitant solution containing 30% w/v PEG 400, 200 mM trisodium citrate (Na^3^C^6^H^5^O^7^) and 100 mM Tris pH 8.5. The crystals were grown by the hanging-drop vapour-diffusion method at 18°C (291 K). Hexagonal crystals of about 0.4 x 0.4 x 0.05 mm^3^ grew within one week. They were mounted free standing supported by a film of buffer in nylon loops (Hampton Research), vitrified and stored in liquid nitrogen. Diffraction was initially verified using the in-house X-ray source (X8Proteum system, Bruker AXS) while maintaining the crystals at 100 K in a nitrogen gas stream (Oxford CryoSystems Ltd, UK).

#### Synchrotron data collection and 3D structure determination

Diffraction data were collected at 100 K and a wavelength = 0.9677 Å (12.812 keV) at beam-line ID30A-3 (or MASSIF-3) of the European Synchrotron Facility (ESRF) in Grenoble, France. During data collection, the crystal was rotated by 0.1° per exposure and 3600 images were collected. The X-ray diffraction images were processed with XDS^3^. The integrated X-ray intensities were analyzed with the program Pointless^4^ to determine the space group symmetry and then scaled with the program Scala^5^ both of which are part of the CCP4 program suite^6,7^.

Using the SAD data, experimental phases were determined, which lead to the calculation of an interpretable electron density map. An initial structural model of sHcp comprising amino acids A2 to K129 was built into the electron density. Iterative rounds of manual model building with Coot^8^ (version 0.9.8.1 EL) and sHcp structure refinement against data extending to 2.05 Å with the software packages REFMAC5^9^ and Phenix^10^. Final refinement as carried out using phenix.refine^11^ of the Phenix program package until the R-factor and R-free factor converged. Structure validation was performed with the web services provided by the wwPDB (https://validate-rcsb-1.wwpdb.org/). The data collection and refinement statistics are summarized in supplementary Table 1. The atomic coordinates and structure factors for sHcp have been deposited with the Worldwide Protein Data Bank (wwPDB) under PDB code 9QWL.

**Table S1.** Data collection and refinement statistics.

| sHcp_SeM* |  |
| --- | --- |
| Wavelength | 0.9677 |
| Resolution range | 22.44 - 1.95 (2.02 - 1.95) |
| Space group | P 6 |
| Unit cell | 77.366 77.366 47.635 90 90 120 |
| Total reflections | 248578 (26002) |
| Unique reflections | 11983 (1189) |
| Multiplicity | 20.7 (21.9) |
| Completeness (%) | 99.64 (99.58) |
| Mean I/sigma(I) | 35.10 (3.28) |
| Wilson B-factor | 46.84 |
| R-merge | 0.05516 (1.127) |
| R-meas | 0.05659 (1.154) |
| R-pim | 0.01246 (0.246) |
| CC1/2 | 1 (0.943) |
| CC* | 1 (0.985) |
| Reflections used in refinement | 11961 (1185) |
| Reflections used for R-free | 1192 (120) |
| R-work | 0.2092 (0.4084) |
| R-free | 0.2331 (0.3649) |
| CC(work) | 0.946 (0.812) |
| CC(free) | 0.960 (0.682) |
| Number of non-hydrogen atoms | 1034 |
| macromolecules | 1015 |
| ligands | 34 |
| solvent | 5 |
| Protein residues | 129 |
| RMS(bonds) | 0.017 |
| RMS(angles) | 1.80 |
| Ramachandran favored (%) | 88.98 |
| Ramachandran allowed (%) | 6.30 |
| Ramachandran outliers (%) | 4.72 |
| Rotamer outliers (%) | 1.79 |
| Clashscore | 3.44 |
| Average B-factor | 67.34 |
| macromolecules | 67.27 |
| ligands | 76.43 |
| solvent | 55.49 |
| *Statistics for the highest-resolution shell are shown in parentheses. |  |

>sHcp M1-K129+ASGGGHHHHHH, 129+11=140 aa., structural Hemolysin-coregulated-protein, based on Q5LDT1, *Bacteroides fragilis* strain NCTC9343 170518 MAFRATLSFAGKEFDVLDCTYSLKRDVDSKGRPSSNIYGGQIRLHVSTDDTSILENMTN QFKPHSGSIVFKKGDEEAKMKELTWENGYITEFTENIDIVGSQPMTITFVVSQAQVIKIGG AQFEQNWPkasGGGHHHHHHH

---

**Figure S1.**
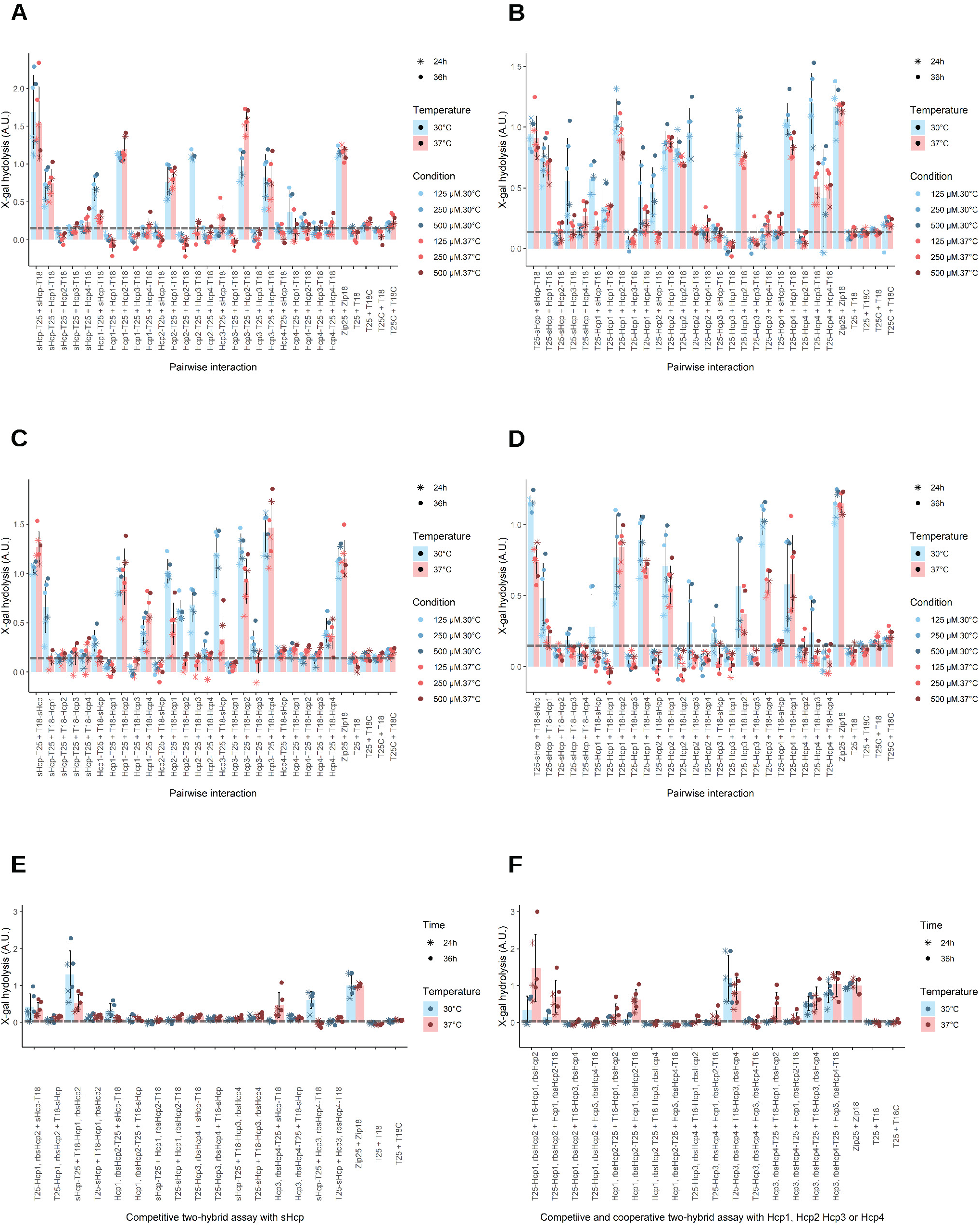
Quantification of normalized X-gal hydrolisis. Quantification of two-hybrid assays of the major and minor Hcp proteins from *B. fragilis* with plasmids carrying genes encoding Hcp proteins fused to either **A)** a C-terminal T18 and T25 fragments or **B)** a N-terminal T25 and a C-terminal T18 fragment or **C)** a C-terminal T25 and a N-terminal T18 fragment or **D)** a N-terminal T18 and T25 fragments. **E)** Quantification of competitive two-hybrid assay with plasmids encoding sHcp fused to N- or C-terminal T18 or T25 fragments tested for pairwise interactions with T18 or T25 fusions of Hcp1, Hcp2, Hcp3 or Hcp4 in the presence of Hcp3, Hcp2, Hcp5 and Hcp4, respectively. **F)** Quantification of competitive and cooperative two-hybrid assay for pairwise interactions pairwise interactions among T18 or T25 translational fusions of Hcp1, Hcp2, Hcp3 and Hcp4 in the presence of Hcp2, Hcp1, Hcp4 and Hcp3, respectively. A dashed line is shown at the average of negative controls.

**Figure S2.**
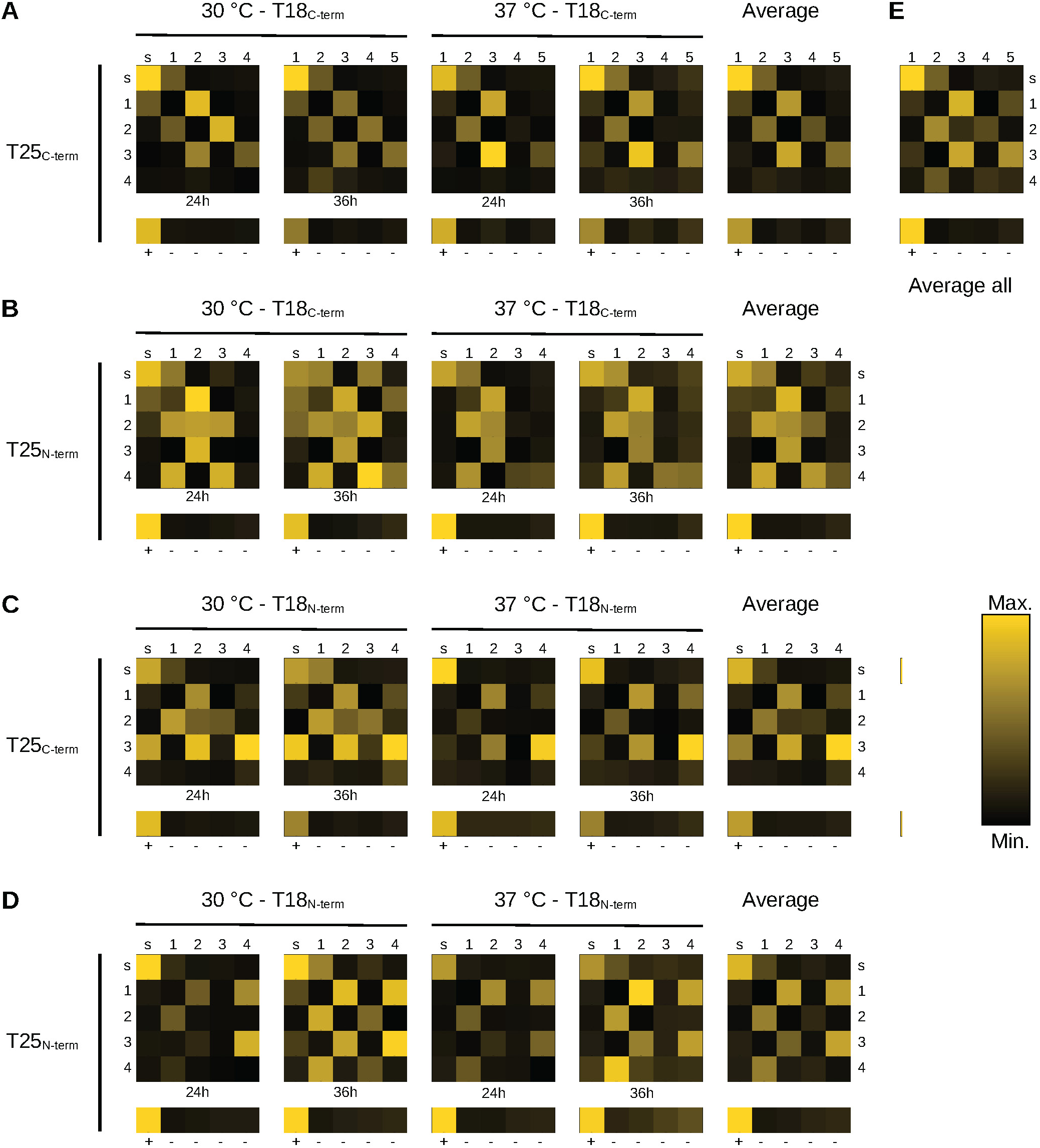
Quantification of two-hybrid assays. The average hydrolized X-gal for each condition tested for two-hybrid assays with plasmids carrying genes encoding Hcp proteins fused to either **A)** a C-terminal T18 and T25 fragments or **B)** a N-terminal T25 and a C-terminal T18 fragment or **C)** a C-terminal T25 and a N-terminal T18 fragment or **D)** a N-terminal T18 and T25 fragments. Hcp proteins are labeled with characters: sHcp (s), Hcp1 (1), Hcp2 (2), Hcp3 (3) and Hcp4 (4). The positive control (+) is the dimerization of the yeast GCN4 leucine zipper and the negative controls (-) are the empty vectors in four different combinations.The data is plotted as a color scale in arbitrary units normalized to both the positive and negative controls. The color scale between black and yellow shows the minimum and maximum, respectively in each individual plot.

**Figure S3.**
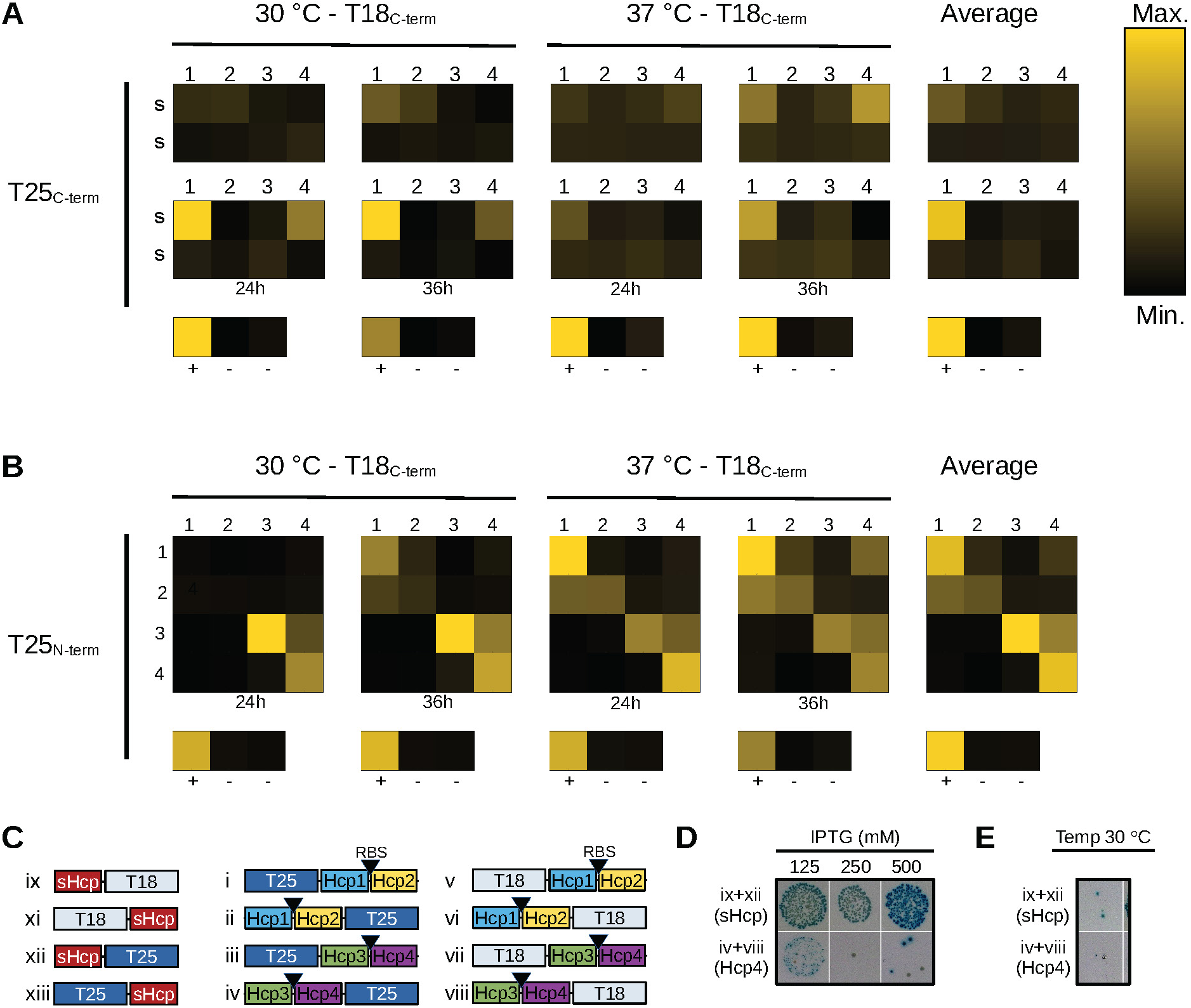
**A)** The average hydrolized X-gal for each condition tested for competitive two-hybrid assay with plasmids encoding sHcp fused to N- or C-terminal T18 or T25 fragments tested for pairwise interactions against T18 or T25 fusions of Hcp1 (1), Hcp2 (2), Hcp3 (3) and Hcp4 (4) in the presence of Hcp2, Hcp1, Hcp4 and Hcp3, respectively. The color scale between black and yellow shows the minimum and maximum, respectively in each individual plot. **B)** The average hydrolized X-gal for each condition tested in competitive and cooperative two-hybrid assay for pairwise interactions between T18 or T25 translational fusions of Hcp1 (1), Hcp2 (2), Hcp3 (3) and Hcp4 (4) in the presence of Hcp2, Hcp1, Hcp4 and Hcp3, respectively. Color scale as in A). **C)** Construct of translational T18 or T25 fusions used for the assay shown in A-B) and they follow the same scheme shown in Figure 1. **D-E)** Effect of IPTG concentration and temperature on two hybrid assays of interaction between sHcp C-terminal translational fusions and between constructs carrying Hcp4 T18 and T25 translational fusions in the rpesence of Hcp3. **D**) Photographic images of two-hybrid assays performed at 37 °C and at different IPTG concentrations. **E)** Photographic images of two-hybrid assays performed at 30 °C at a concentration of 125 mM IPTG.

**Figure S4.**
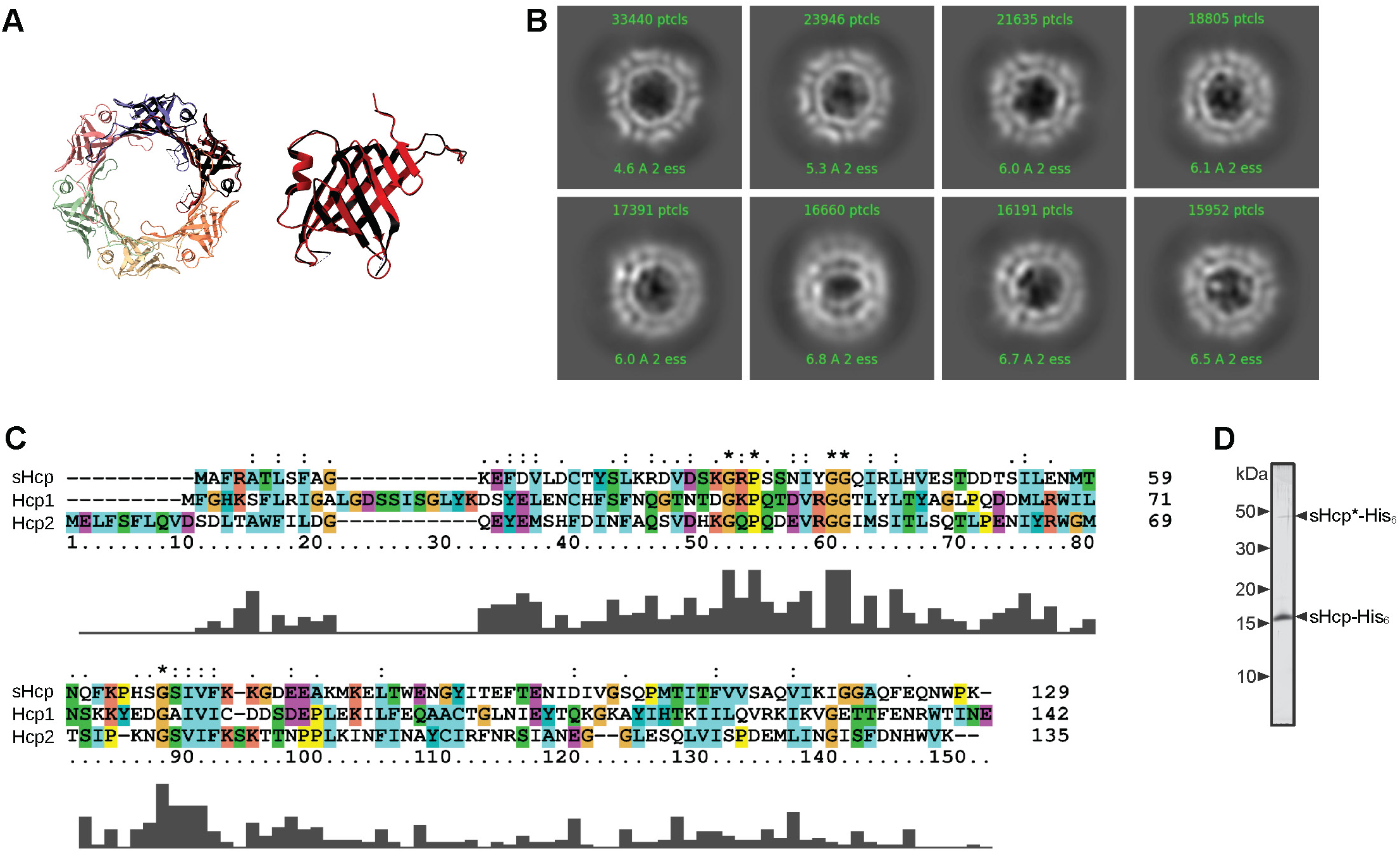
**A)** Structural alignment of to the He *et al.* unbound structure (black) to our sHcp hexameric crustal structure (left) and monomer (right). **B)** Top six 2-D classes averaged from Cryo-EM data from Hcp1-Hcp2-His_6_. **C)** Chimera structural sequence alignment of sHcp with Hcp1 and Hcp2. **D)** Phosphoimager scan of SYPRO Ruby stained SDS-PAGE gels of purified sHcp-His_6_ at higher exposure.

**Figure S5.**
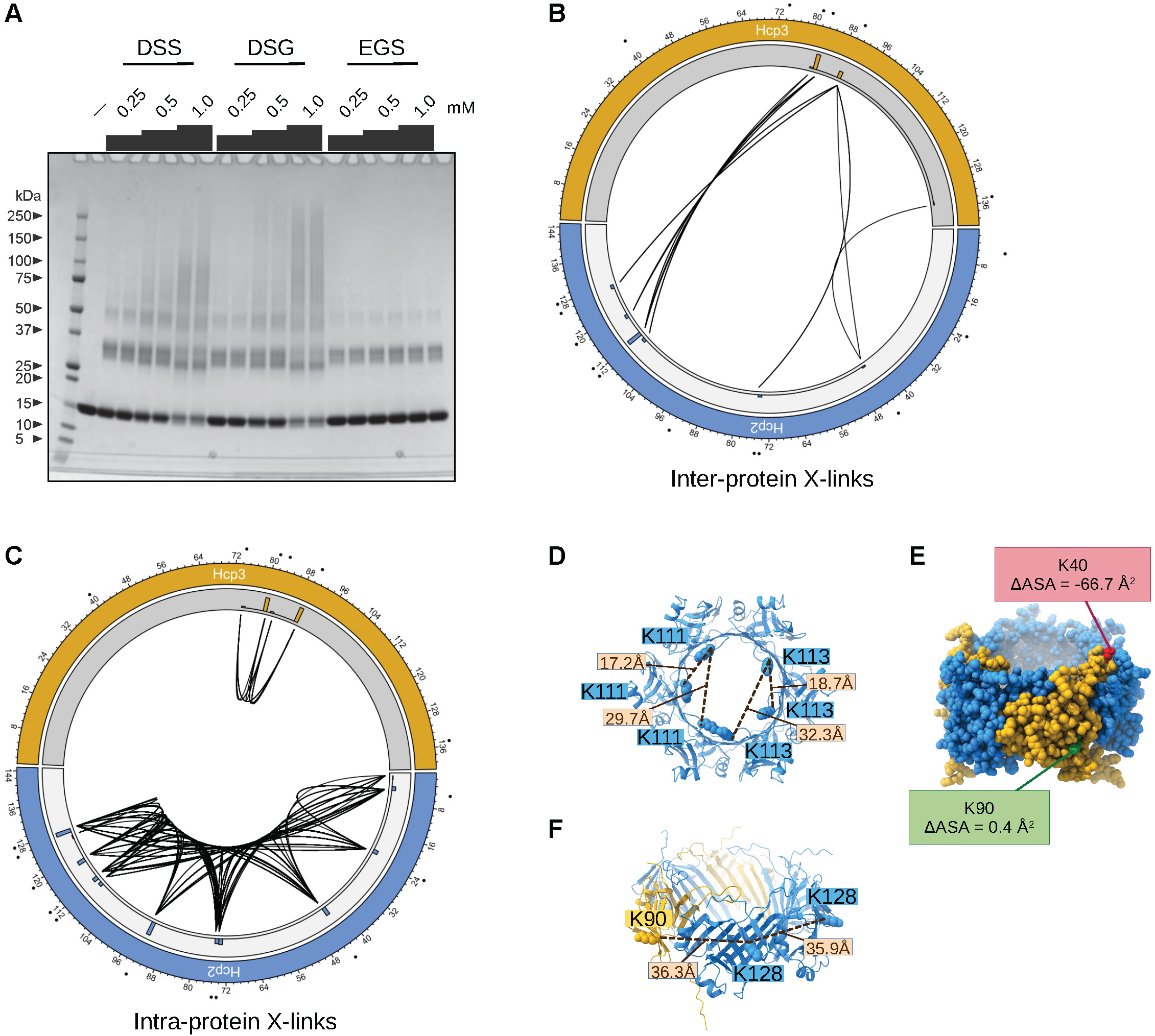
Crosslinking MS of the Hcp1-Hcp2 complex show homo-Hcp1 interactions. **A)** SDS-PAGE of X-linked Hcp proteins using DSS, DSG and EGS. **B-C)** Circos plot of all x-linked peptides found in inter- and intra-protein X-links. Gray track contains a histogram of the total number of peptides found for each residue. **D)** 4:2 complex showing K40 in Hcp2, which was not found in any X-link. E) 4:2 complex showing lysines in Hcp1 involved in interprotein X-links as determined by their X-linking to the same residue in another Hcp1 protein.

**Figure S6.**
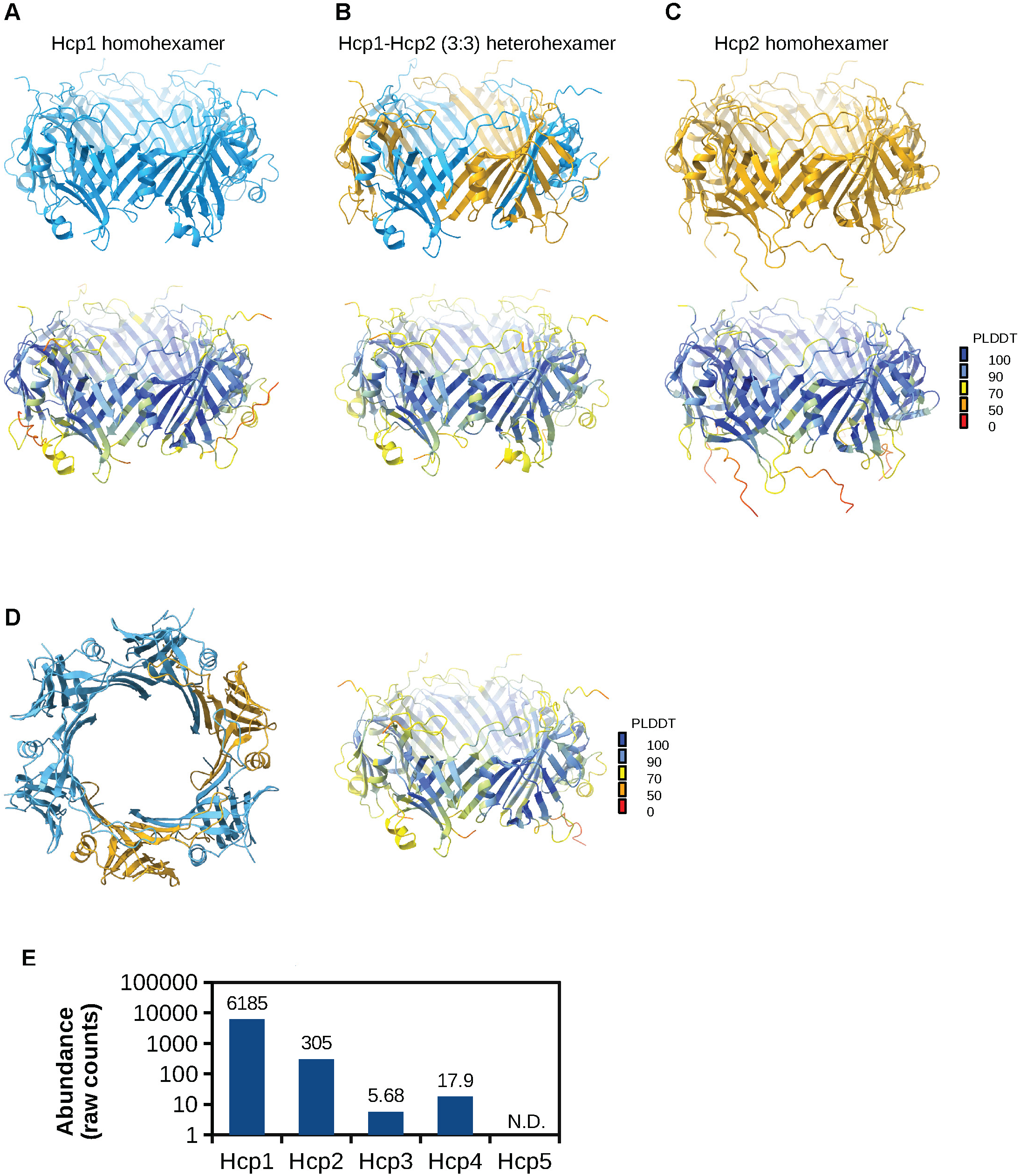
AlphaFold server models of Hcp1 and Hcp2. **A)** Homohexameric model of Hcp1 with PLDDT score(bottom). **B)** Heterohexameric model of Hcp1 and Hcp2 with PLDDT colour-coded structure (bottom) in a 3:3 ratio. **C)** Homohexameric model of Hcp2 with PLDDT colour-coded structure (bottom). **D)** AlphaFold model of a Hcp1-Hcp2 complex in a 4:2 complex with PLDDT colour-coded structure (right side). Hcp1 shown in blue and Hcp2 shown in yellow. **E)** Abundance of Hcp proteins in *B. fragilis* according to Müller, *et al.* (https://doi.org/10.1038/s41586-020-2402-x).

**Figure S7.**
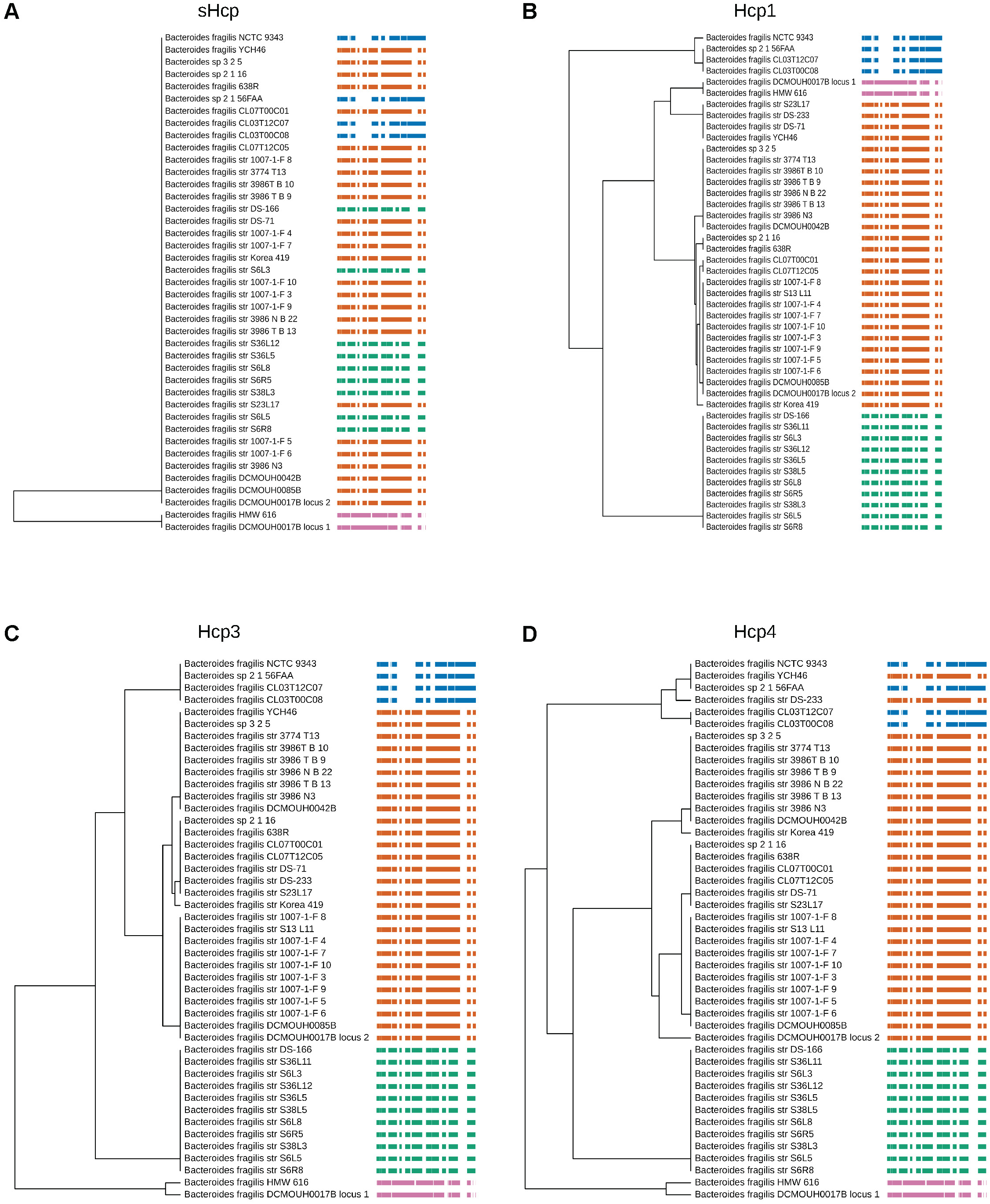
Clustering of phylogenetic trees of sHcp, Hcp1, Hcp3 and Hcp4 proteins from GA3 T6SS with effector proteins. Bte1 (blue), Bfe1 (orange), Cluster 51 (green), Cluster 115 (magenta). Generated with TreeViewer. Trees were compressed horizontally.

**Figure S8.**
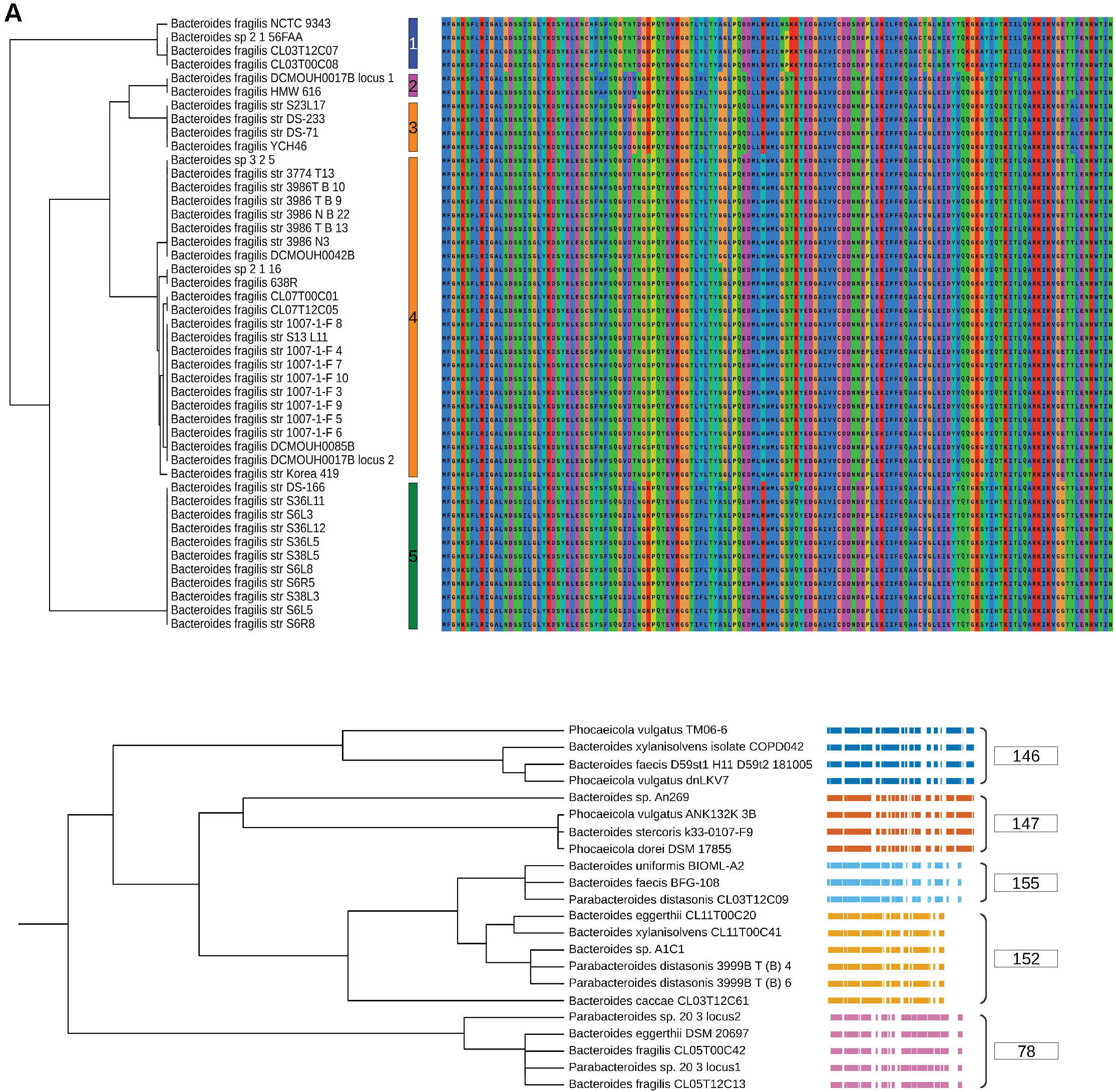
Hcp2 determines binding to Bacteroidales GA2 and GA3 T6SS effectors. **A)** Clustering of phylogenetic tree of Hcp2 proteins from GA3 T6SS with their corresponding sequence coloured with Clustal scheme. **B)** Clustering of phylogenetic trees of Hcp2 proteins from Bacteroidales T6SS GA2 architecture with their effectors (colour bars). The cluster number (78, 146, 147, 152 and 155) designates the HMM model named by Coyne, *et al.* 2016 (16). Effectors were aligned with the MAFFT-G-INS-1 algorithm and clustered by average (UPGMA) distance in JalView before adding the different aligned cluster sequences (coloured bars) to the phylogenetic tree. Spaces between the coloured bars (sequences) are gaps in the global alignment of all effectors. Generated with TreeViewer.

